# Sample Preparation and Homology-Driven Search Shape the First Holobiont Proteome of the Marine Sponge *Viles ophiraphidites* (formerly *Topsentia ophiraphidites*)

**DOI:** 10.64898/2026.10.20.760564

**Authors:** Carolaine da Silva de Araújo, Ana Caroline Silas Rodrigues, Gabrielle Rodrigues de Souza, Fábio César Souza Nogueira, Alessandra Leda Valverde, Magno Rodrigues Junqueira

## Abstract

Marine sponges are holobionts of host cells, extracellular matrix and dense microbial communities, and most lack reference proteomes. *Viles ophiraphidites* (formerly *Topsentia ophiraphidites*), a Brazilian demosponge, is represented in UniProtKB by a single protein. The study had two phases. In phase 1, four sample-preparation workflows were compared on two specimens in 42 nLC-MS/MS runs: urea/thiourea in-solution digestion, acetone precipitation, S-Trap and GeLC-MS/MS. Every run was analysed with FragPipe and with a homology-driven pipeline that adds three layers to a rescored Sage search: the proteome of the closest sequenced relative (*Halichondria panicea*), MS-BLAST-style alignment of de novo sequence tags, and error-tolerant interpretation of tag flanks. In phase 2, the selected workflow, S-Trap, was applied to four specimens in 12 runs. Both strategies ranked the workflows identically (ρ = 0.98). S-Trap gave the deepest and most specimen-independent proteome, with about 2,100 protein groups per specimen and an 8% difference between specimens, against 21–39% for the other workflows. The homology-driven layers raised assigned non-contaminant spectra by 21.3% in phase 1 and 26.8% in phase 2. They contributed 1,969 protein groups found only by homology and agreed with the database search on 99.5–99.9% of shared spectra. The proteome comprises 4,294 protein groups: 42% sponge and 58% bacterial or archaeal, the latter dominated by Poribacteria. It includes *Nitrospira* nitrite oxidoreductase detected in every run. The four phase-2 specimens had a similar composition (Bacteria 51–64% and Porifera 35–48% of peptides, Poribacteria alone 27–33%), whereas the two specimens stored long-term in ethanol at 4 °C, which also differed in collection site and storage history, yielded four to five times less evidence than the frozen ones, even though reduction and alkylation had not been effective in the two frozen specimens. S-Trap combined with homology-driven search is a robust strategy for proteomics of non-model holobionts.

## INTRODUCTION

Most eukaryotic species still lack a genome, and fewer still have experimentally supported proteomes. Fewer than 1% of described eukaryotes have a representative genome in GenBank^1^. For such non-model organisms, tandem mass spectra must be searched against sequences of related taxa. Every amino-acid difference between the organism and its database relatives then costs an identification^2^.

Marine sponges show why this gap matters. They filter large volumes of seawater, transfer dissolved organic matter to benthic food webs through the sponge loop, and host dense microbial communities that drive carbon, nitrogen and phosphorus cycling^3−5^. A sponge sample is a holobiont: host cells, a collagenous and spicule-rich extracellular matrix, and bacteria and archaea whose peptides share the same spectra. Proteomics of this mixture must recover proteins from very different cell types. It must also assign each peptide to host or symbiont against databases in which neither is well represented.

*Viles ophiraphidites* (de Laubenfels, 1934) is common in the tropical western Atlantic and on oceanic islands of Brazil, including Fernando de Noronha^6^. Long treated as *Topsentia ophiraphidites* (Halichondriidae), it is now the type species of the resurrected genus *Viles*, placed in the new family Vilesidae and order Vilesida^7^. In October 2026 UniProtKB held a single unreviewed protein for this species (mitochondrial COX1)^8^, and NCBI Gene listed 41 records^9^. The closest relative with an annotated genome is *Halichondria panicea* (Halichondriidae), from the family in which the species was classified until 2025, with a RefSeq-annotated assembly (GCF_963675165.1, odHalPani1.1). A first protein-level description of *V. ophiraphidites* therefore depends on two choices made before and after the mass spectrometer.

The first choice is sample preparation. Tissue disruption, solubilisation, detergent removal and digestion determine which proteins reach the instrument^10,11^. Chaotrope-based in-solution digestion, acetone precipitation, suspension-trap (S-Trap) digestion and in-gel digestion differ in how they handle SDS, matrix and salts. No workflow has been evaluated for this species or its close relatives.

The second choice is how spectra are interpreted. A conventional database search reports only peptides whose exact sequence is in the database. Homology-driven proteomics adds the peptides of proteins that differ from their database relatives. In this approach, de novo sequence candidates are aligned to related proteins with mass-spectrometry-aware substitution matrices, as in MS-BLAST^12^. Error-tolerant sequence-tag searching^13^ extends this idea. A confidently read stretch of the spectrum anchors the peptide, and the unexplained mass is attributed to substitutions, non-tryptic ends or modifications. Modern rescoring of database-search results with semi-supervised machine learning (Percolator, mokapot) also raises the number of PSMs accepted at a fixed FDR^14,15^.

Here we studied *V. ophiraphidites* in two phases (Figure 1). In phase 1, we analysed 42 LC-MS/MS runs from two specimens prepared by four workflows: urea/thiourea in-solution digestion (A-UT), acetone precipitation (A-PPT), S-Trap (BST) and GeLC-MS/MS (BGEL). In phase 2, we applied the best workflow, S-Trap, to four specimens (12 runs) to test whether the proteome and the holobiont composition hold across individuals. Every run was processed with two independent strategies: FragPipe (MSFragger, MSBooster, Percolator, IonQuant), a widely used reference workflow, and the LEMTEN pipeline, which adds de novo sequencing, homology search and error-tolerant sequence tags to a rescored Sage search against a database extended with the *H. panicea* proteome.

**Figure 1.**
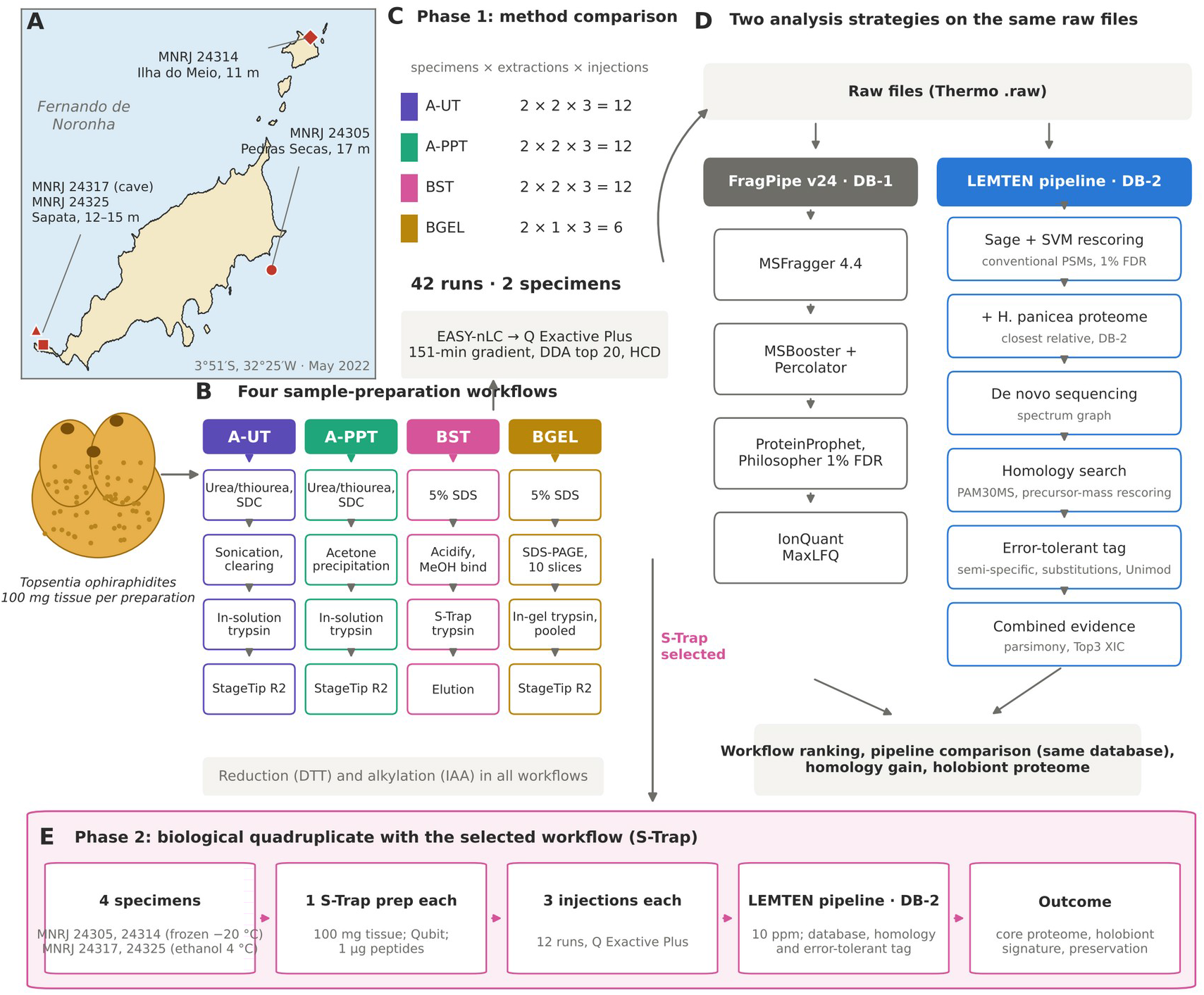
Two-phase study design and data analysis. (A) Collection sites at Fernando de Noronha (May 2022): MNRJ 24305 (Pedras Secas, 17 m) and MNRJ 24314 (Ilha do Meio, 11 m), used in both phases; MNRJ 24317 (Caverna da Sapata, 12 m) and MNRJ 24325 (Ponta da Sapata, 12–15 m), used in phase 2. Schematic of the sponge (100 mg tissue per preparation). (B) The four sample-preparation workflows of phase 1 and their main steps: A-UT, A-PPT, BST and BGEL. (C) Phase 1 replicate structure: two extractions × three injections for A-UT, A-PPT and BST, and one pooled gel lane × three injections for BGEL; 42 runs. (D) The two analysis strategies: FragPipe on DB-1, and the LEMTEN pipeline on DB-2 (DB-1 plus *Halichondria panicea*). LEMTEN steps: Sage with SVM rescoring, de novo sequencing, PAM30MS homology search with precursor-mass and multivariate rescoring, error-tolerant sequence tag, combined protein inference. (E) Phase 2: the selected workflow (S-Trap) applied to four specimens, two kept frozen at −20 °C and two stored in 80% ethanol at 4 °C, each injected three times (12 runs) and analysed with the LEMTEN pipeline.

**Figure 2.**
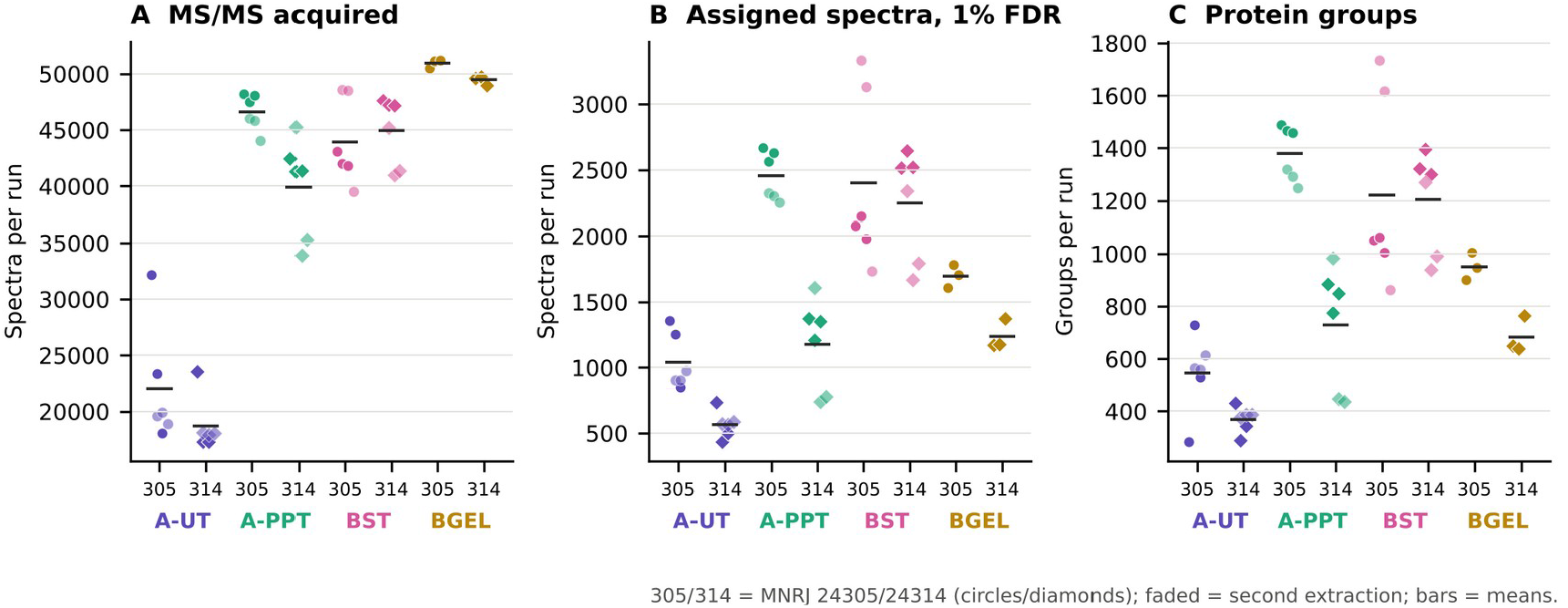
Identifications per workflow and specimen. Per-run MS/MS spectra, assigned spectra and protein groups (LEMTEN pipeline); points are individual injections, coloured by extraction.

We asked four questions. First, which workflow recovers the most proteome, most reproducibly, in both specimens? Second, how do the two search strategies agree, and where do they differ? Third, what do PSM, homology and sequence-tag evidence together reveal about the first proteome of this sponge holobiont? Fourth, is this proteome reproducible across specimens with the selected workflow? The answers define a sample-preparation and search strategy for sponge holobionts without reference proteomes, and provide a first proteome of the species.

## EXPERIMENTAL PROCEDURES

### Specimens

Two specimens of *Viles ophiraphidites* (de Laubenfels, 1934), MNRJ 24305 and MNRJ 24314, were collected at the Fernando de Noronha Archipelago in May 2022 by the team of Dr. Guilherme Muricy (Museu Nacional, UFRJ). MNRJ 24305 was collected off Pedras Secas Island (17 m) and MNRJ 24314 at Carvinhas, Ilha do Meio (11 m). Both specimens were kept frozen at −20 °C in the laboratory. Collection is registered in SisGen (AB724BB).

### Protein Extraction

For each workflow, 100 mg of tissue was homogenised with a microtube pestle in one of two lysis buffers, both containing cOmplete Mini EDTA-free protease inhibitor (Roche): buffer A (7 M urea, 2 M thiourea, 0.5% (w/v) sodium deoxycholate, 50 mM TEAB, pH 8.5) or buffer B (5% SDS, 50 mM TEAB, pH 8.5). The workflows that used each buffer are described below.

#### Urea/thiourea in-solution digestion (A-UT)

Lysates in buffer A were sonicated (Ecosonics; 15 cycles of 10 s on/10 s off, 30% power) and cleared at 16,000 × g for 10 min at 4 °C. Proteins (200 µg) were reduced with 10 mM DTT (1 h, 37 °C) and alkylated with 40 mM iodoacetamide (30 min, dark).

#### Acetone precipitation (A-PPT)

An aliquot of the A-UT extract was precipitated overnight with ice-cold acetone at −20 °C. The pellet was washed three times, alternating ice-cold acetone and methanol, with centrifugation at 10,000 × g and 4 °C after each wash. It was then dried in a SpeedVac and resolubilised in 8 M urea, 100 mM TEAB. Reduction and alkylation were as for A-UT.

#### S-Trap (BST)

Lysates in buffer B were sonicated, cleared, reduced and alkylated as above. They were acidified to 1.2% phosphoric acid, diluted 1:7 in binding buffer (90% methanol, 100 mM TEAB, pH 7.5) and loaded on S-Trap micro columns (ProtiFi)^16^. After three washes, proteins were digested with trypsin (1:50, 125 µL in 50 mM TEAB) for 18 h at 37 °C. Peptides were eluted sequentially with 50 mM TEAB, 0.2% formic acid and 50% acetonitrile.

#### GeLC-MS/MS (BGEL)

About 200 µg of the buffer-B extract was separated on a 15% SDS-PAGE mini-gel at 200 V for 90 min and stained with Coomassie G-250. One lane per specimen was excised into 10 slices and subjected to in-gel digestion^17^. Gel slices were destained with 100 mM ammonium bicarbonate/acetonitrile (1:1), reduced with 10 mM dithiothreitol (DTT) and alkylated with 40 mM iodoacetamide. After dehydration with acetonitrile, proteins were digested with 12.5 ng/µL trypsin in 50 mM ammonium bicarbonate for 18 h at 37 °C. Peptides were extracted with 5% formic acid/acetonitrile (1:2). Peptide extracts from all 10 slices of each lane were pooled into one sample per specimen, and each pooled sample was analysed in three technical LC-MS/MS injections.

### Digestion, Cleanup and Quantification

A-UT and A-PPT proteins (200 µg) were diluted tenfold in 50 mM TEAB and digested with trypsin (Promega; 1:50, 18 h, 37 °C). Digestion was stopped with 1% formic acid. A-UT, A-PPT and BGEL peptides were desalted on in-house Poros R2 StageTips: conditioning in acetonitrile, equilibration and washes in 0.1% TFA, elution in 50% and 70% acetonitrile with 0.1% TFA. Proteins and peptides were quantified with a Qubit 2.0 fluorometer (Invitrogen).

### Experimental Design

#### Phase 1 (workflow comparison)

A-UT, A-PPT and BST were each prepared as two independent extractions per specimen; BGEL as one preparation per specimen. Each peptide sample was injected three times, giving 42 runs: 12 A-UT, 12 A-PPT and 12 BST runs (2 specimens × 2 extractions × 3 injections) and 6 BGEL runs (2 specimens × 1 preparation × 3 injections).

Blank and quality-control runs from the same series were not analysed. Injections are technical replicates and extractions measure preparation repeatability. The two specimens are the only biological replicates of phase 1, so comparisons between workflows are descriptive.

#### Phase 2 (biological quadruplicate)

The workflow selected in phase 1, S-Trap, was applied to four specimens, MNRJ 24305, 24314, 24317 and 24325. Each specimen gave one preparation, injected three times, for 12 runs (see Phase 2: Biological Quadruplicate below).

### nLC-MS/MS

Peptides (1 µg) were analysed on an EASY-nLC system coupled to a Q Exactive Plus mass spectrometer (Thermo Fisher Scientific). Peptides were separated on an EASY-Spray PepMap RSLC C18 column (75 µm × 50 cm, 2 µm particle size, 100 Å pore size) at 300 nL/min. Mobile phases A and B were 5% and 95% acetonitrile, respectively, each with 0.1% formic acid. The gradient started at 3% B and increased to 23% B at 81 min, 35% B at 111 min, 45% B at 121 min and 97% B at 131 min; this composition was held until 141 min, and the gradient returned to 3% B at 151 min.

Spectra were acquired in data-dependent mode with a top-20 method. Full MS scans covered m/z 350–2,000 at a resolution of 70,000, with an automatic gain control (AGC) target of 3 × 10⁶ and a maximum injection time of 100 ms. MS/MS spectra were acquired after higher-energy collisional dissociation (HCD) at a normalised collision energy of 30, with an isolation window of 1.2 m/z, a resolution of 17,500, an AGC target of 1 × 10⁶ and a maximum injection time of 50 ms. The precursor intensity threshold was 2 × 10⁵, singly charged precursors and those with unassigned charge were excluded, and dynamic exclusion was set to 45 s.

### Protein Sequence Databases

The laboratory database combines UniProtKB entries for Porifera and sponge-associated microbial groups: Actinobacteria, Alphaproteobacteria, Archaea, Bacteroidota, Chloroflexi, Cyanobacteria, Gammaproteobacteria, Nitrospirota, Planctomycetota, Poribacteria, Proteobacteria and Verrucomicrobiota. Files were concatenated in Galaxy and made non-redundant with PatternLab V^18^, giving 473,966 sequences (DB-1).

For the LEMTEN pipeline, DB-1 was extended with the RefSeq proteome of *Halichondria panicea* (GCF_963675165.1, odHalPani1.1; 23,028 non-redundant proteins). This species is the closest relative of *V. ophiraphidites* with an annotated genome. The extended database (DB-2) held 497,112 targets plus 118 contaminants, searched with an equal number of reversed decoys.

### FragPipe Workflow

Raw files were searched against DB-1 with MSFragger^19^ v4.4.1 within FragPipe v24.0. Searches used full tryptic specificity with up to two missed cleavages, and precursor and fragment mass tolerances of ±10 ppm and 0.02 Da, respectively. Carbamidomethylation of cysteine was a fixed modification, and methionine oxidation and protein N-terminal acetylation were variable modifications, with at most three variable modifications per peptide. MSBooster^20^ v1.4.14 generated predicted fragment intensities and retention times for rescoring with Percolator^14^ v3.7.1. ProteinProphet performed protein inference, and PSMs, peptides and proteins were filtered at 1% FDR with Philosopher^21^ v5.1.3. IonQuant^22^ provided MaxLFQ intensities across the 42 runs.

### LEMTEN Pipeline

Each raw file was processed independently (pipeline version of 11 October 2026) in the eight steps described below.

#### Conversion and database search

ThermoRawFileParser^23^ produced MGF and mzML files. Sage^24^ v0.14.7 searched DB-2 in eight database chunks, with trypsin specificity, up to two missed cleavages and peptide lengths of 7–30 residues. Carbamidomethylation of cysteine was a fixed modification and methionine oxidation a variable modification (at most two per peptide). Precursor and fragment tolerances were ±20 ppm, with precursor isotope errors from −1 to +2, and chimeric-spectrum search and retention-time prediction were enabled. The best PSM per spectrum across chunks was rescored by a semi-supervised linear support vector machine (SVM), in the style of Percolator and mokapot^15^. The model used 17 Sage scores and the deviation from the predicted retention time as features and was trained with three-fold cross-validation by spectrum over three iterations. PSM and peptide FDR were set at 1% by target–decoy competition, with picked-protein FDR and parsimony grouping at the protein level.

#### Label-free quantification

Peptide abundance was the MS1 extracted-ion chromatogram area (M0+M1+M2, ±10 ppm). Protein abundance was the mean of the three most intense peptides (Top3). Match-between-runs transferred areas of non-fragmented peptides for quantification only.

#### De novo sequencing

A spectrum-graph algorithm scored b/y prefix-mass nodes and found the best path to the precursor mass by dynamic programming, also testing M − 1.00335 Da. In every run, accuracy was measured against the 1% FDR PSMs as amino-acid precision and peptide recall.

#### Homology search

De novo candidates were aligned to DB-2 targets and decoys with a PAM30MS matrix (MS-BLAST style). An alignment gained a bonus when the tryptic database peptide containing it matched the precursor mass (≤15 ppm; Cys alkylated; ≤2 oxidised Met; isotope 0/+1). A smaller bonus applied when the residual mass equalled one amino-acid substitution.

#### Error-tolerant sequence tag

The aligned stretch was treated as a sequence tag^13^. The pipeline then tested two explanations for the unaligned flanks. The first was a semi-specific peptide, with up to four flanking residues lost to non-tryptic cleavage or in-source fragmentation. The second was a modification among the 1,543 in Unimod^25^, matched within ±0.01 Da (|Δ| ≥ 2.1 Da), whose site lay in an unaligned flank; this explanation was used for annotation only.

Precursor–peptide mass differences of hits at q ≤ 0.05 were binned at 0.01 Da, with target and decoy hits in parallel. Each recurrent bin was annotated as an amino-acid substitution, a Unimod modification or an isotope error.

#### Rescoring and FDR of homology hits

A linear discriminant separated the best target from the best decoy alignment per spectrum, with three-fold cross-validation. Its features were the alignment score, identities, alignment length and identity fraction; the query length, coverage and fraction of unassigned residues (X); mass agreement, substitution and |ppm|; and the de novo score, number of residues, charge and semi-specific flag.

Spectrum-level target–decoy competition set a 1% FDR. Spectra already identified by Sage served as a positive control of the observed error.

#### Combined evidence and comparison

For spectra without a Sage PSM, the homology hit was added, and protein groups were rebuilt by parsimony from the combined set. Each group was annotated by identification source (Sage, homology or both) and origin (sponge, symbiont [Bacteria or Archaea] or contaminant). Runs were combined according to the design, averaging injections within each preparation and preparations within each specimen. For each workflow, the pipeline reported identifications, coefficients of variation between injections, extractions and specimens, principal component analysis (PCA) of log₂ Top3 abundances, overlap among workflows, and spectral abundance factor (SAF) rank–abundance curves highlighting the 16 candidate proteins.

Gene Ontology (GO) terms were assigned from the best *H. panicea* homologue using RefSeq GAF annotations, propagated to ancestor terms. Additional terms came from the best human Swiss-Prot homologue found with DIAMOND (E ≤ 10⁻⁵, identity ≥ 30%, coverage ≥ 50%).

### Comparison of the Two Workflows

The outputs of both pipelines were compared run by run after excluding contaminants (*Homo sapiens*, *Mus musculus*, *Sus scrofa* and *Bos taurus*). PSMs were compared as summed FragPipe spectral counts against LEMTEN PSMs at 1% FDR. Protein-level comparisons used FragPipe proteins with at least one unique spectrum and LEMTEN protein groups, and a FragPipe protein was counted as shared when its accession was a member of a LEMTEN group. For two BST runs, FragPipe psm.tsv files were also matched to LEMTEN PSMs scan by scan, treating isoleucine and leucine as equivalent.

### Peptide-Level Taxonomic Assignment

Every distinct non-contaminant peptide accepted at 1% FDR in any of the 42 runs (database and homology search, I and L equated) was mapped to the organisms of all database proteins that contain it, read from the OS= field of UniProt headers; RefSeq entries were assigned to *Halichondria panicea*. Organisms were placed in a domain–phylum–class lineage following the NCBI Taxonomy, and each peptide was assigned to the deepest rank shared by all its proteins, the lowest common ancestor (LCA), as in Unipept^26^. The tree was drawn with lca_tree.py from the LEMTEN pipeline kit.

### Phase 2: Biological Quadruplicate

To test reproducibility across specimens, four specimens were analysed by S-Trap: MNRJ 24305 and MNRJ 24314 (above), and MNRJ 24317 and MNRJ 24325, collected on 9 May 2022 at Fernando de Noronha by Guilherme Muricy and Joana Sandes. MNRJ 24317 came from the second chamber of Caverna da Sapata (12 m) and MNRJ 24325 from Ponta da Sapata, outside the cave, growing within a large *Ectyoplasia* specimen (12–15 m).

Storage differed between the two pairs. MNRJ 24305 and MNRJ 24314 had been kept frozen at −20 °C in our laboratory, whereas MNRJ 24317 and MNRJ 24325, obtained later from the Museu Nacional collection, had been kept in 80% ethanol at 4 °C. To equalise storage before extraction, a piece of each frozen specimen was transferred to 80% ethanol, and all four were held in ethanol at −20 °C for one month. Residual ethanol was then evaporated, and the tissue was washed with PBS and dried; 100 mg of tissue was then cut from each specimen for extraction.

One S-Trap preparation was made from 100 mg of tissue of each specimen, as described for BST. For MNRJ 24305 and MNRJ 24314, the cysteine forms in the data show that reduction and alkylation were not effective in this preparation (see Results); the cause could not be established. Protein and peptide concentrations were measured by Qubit. One microgram of peptides per specimen was injected in technical triplicate, giving 12 runs (QE_018062–QE_018076). The instrument was well calibrated for these runs (median precursor error 1.7–3.1 ppm). They were therefore searched with a 10 ppm precursor and 0.02 Da fragment tolerance, with protein N-terminal acetylation as an additional variable modification and up to three modifications per peptide, against DB-3 (see Database Construction, Optimization and Testing below). All other settings were as for the LEMTEN pipeline above. The same runs were also searched in FragPipe against DB-3 with the phase-1 FragPipe settings.

The runs were compared by specimen (bio) and injection (tec) with the same scripts, without match-between-runs. Peptide-level LCA trees were built for each specimen.

### Cysteine Alkylation Check

Reduction and alkylation were checked in all 54 runs (42 of phase 1 and 12 of phase 2). Each run was searched with Sage against the 6,001 proteins identified in 12 representative runs (both specimens and all workflows of phase 1, one injection per specimen of phase 2) and their reversed decoys. No modification was fixed. Variable modifications on cysteine were carbamidomethylation (+57.0215 Da), disulfide bonding (−1.0078 Da), dioxidation, trioxidation, propionamide, carboxymethyl, N-ethylmaleimide, pyridylethyl, methylthio and two other adducts, with Met oxidation and at most two modifications per peptide. Target PSMs at 1% FDR without contaminant matches were counted, and the forms of cysteine were tallied per run (Supporting Table S9).

### Database Construction, Optimization and Testing in an LLM-Agent Environment

The phase-2 database was audited, optimized and tested with an LLM agent (Claude, Anthropic) working in an isolated Linux workspace that held the LEMTEN pipeline kit, FragPipe v24.0, Sage v0.14.7, MMseqs2 (release 17-b804f) and DIAMOND v2.1.10, as described for our earlier agent-based analyses^27^. The agent ran every step as code and kept the scripts, logs and intermediate files. It proposed each change to the database with the supporting numbers, and the authors decided whether to adopt it.

#### Construction and integrity

DB-2 (above) was the starting point: 473,966 sequences inherited from DB-1, the 23,028 *H. panicea* proteins and 118 contaminants. Every inherited entry was checked against the source FASTA by identifier and sequence, and only two identical sequences were found. Organism names were read from the OS field of the headers rather than from accessions, because many TrEMBL accessions of DB-1 are no longer active in UniProtKB.

#### Audit

DB-2 was examined against the phase-2 identifications obtained with it (FragPipe, 12 runs) in four ways. For composition, the share of database residues of each taxon was compared with its share of razor PSMs. For redundancy, sequences were clustered with MMseqs2 linclust at 95% and 70% identity and coverage 0.8^28^. For taxonomic consistency, the 52,163 entries labelled *Geodia barretti* were aligned to the rest of DB-2 with DIAMOND (blastp, fast mode, five hits per query, E ≤ 10⁻⁵)^29^; an entry was classed as bacterial when its best bacterial or archaeal hit scored at least 1.5 times its best sponge hit, or when no sponge protein was among its hits. Finally, peptide sharing was measured as the number of identified peptides that occur in proteins of more than one taxonomic group.

#### Optimization

Two candidate databases were derived from DB-2. Both replaced the contaminant list with the union of the original 118 entries and the universal contaminant library^30^, giving 429 contaminants, and both kept the *Geodia* entries classed as bacterial, relabelled as unclassified Bacteria for all taxonomic summaries (protein origin and LCA). DB-3a also removed all *Vibrio* and Planctomycetota entries (186,436 sequences, including Phycisphaerae, *Rhodopirellula* and *Scalindua*), which together held 35% of the residues of DB-2 but explained 1.1% of the PSMs. DB-3 removed only the *Vibrio* entries (30,680 sequences), which are culture isolates from other sponges and explained 0.1% of the PSMs. Clustering was not adopted, because it reduced the database by only 8.6% at 95% identity.

#### Testing

Both candidates were tested against DB-2 before adoption. First, one injection per phase-2 specimen was searched with Sage against each database extended with an entrapment set: about half of the target proteins, with the residues of every tryptic peptide shuffled and its C-terminal K or R kept in place, plus reversed decoys of targets and entrapment sequences. The false discovery proportion among accepted PSMs was estimated with the combined method, FDP = N_e(1 + 1/r)/(N_t + N_e), where N_t and N_e are accepted target and entrapment PSMs and r is the entrapment-to-target size ratio, here 0.50^31^. Second, the 12 phase-2 runs were re-searched in FragPipe with the workflow above, changing only the database.

The FragPipe test exposed a pitfall worth reporting. When decoys were first generated with headers carrying only the accession, MSFragger’s protein-evidence grouping (group_variable = 2, which reads the PE field of UniProt headers) put every decoy in one group and most targets in others. Percolator learned the group as a target label, and one run returned 14,127 PSMs at 1% FDR, 2.5 times the DB-2 result; the additional PSMs had a median hyperscore of 13, against 26 for the others. Regenerating the decoys with the full target header, prefixed by rev_, made the target and decoy PE distributions identical and removed the excess. Databases assembled outside FragPipe should therefore keep the full header in their decoys.

A second error was found in the LEMTEN pipeline itself, while the agent investigated why homology-only hits fell with DB-3. The rescoring step of the homology layer rebuilt each decoy protein as the full reverse of its target, whereas the decoys of DB-1 and DB-2 keep the initial methionine and reverse the rest of the sequence. The decoy peptides used to calibrate the homology layer therefore differed from those that had been searched, and its target-decoy threshold was too permissive. The step now reads the decoy sequences from the searched FASTA. The error affected the searches on DB-1 and DB-2. The DB-3 decoys are full reverses, so the earlier step had reconstructed them correctly; reprocessing one phase-2 run with the corrected step gave an identical result. The 42 phase-1 runs were reprocessed with the corrected step; their database-search PSMs were unchanged, the homology layer reported 31% fewer new spectra, and the observed error among positive-control spectra at a nominal 1% FDR fell from 2.0% to 1.0% (Results).

#### Adoption

DB-3a gave the largest gain in identifications, but it removed the Planctomycetota from the analysis although their peptides made up 2–5% of the LCA-assigned peptides of every phase-2 specimen with DB-2, including the anammox class Brocadiia. DB-3 keeps them and was adopted for all phase-2 searches, both LEMTEN and FragPipe (466,314 targets and 429 contaminants; MD5 c392944394ee74f2448fa0bacdff7943). Phase 1 was not re-searched. Results of all three databases are in Supporting Table S10.

## RESULTS AND DISCUSSION

### Identification Performance of the Four Workflows

Across the 42 runs, 1,598,126 MS/MS spectra were acquired. The LEMTEN pipeline assigned 76,936 of them at 1% FDR. Of these, 8,786 (11%) matched only contaminants, almost all keratins and trypsin in the gel runs, leaving 68,150 sponge or symbiont spectra. A-PPT and BST gave the deepest coverage in both specimens; A-UT gave the shallowest. The two specimens differed more than any two replicate extractions. MNRJ 24314 yielded 29–52% fewer PSMs than MNRJ 24305 with every workflow except BST (−8%).

The MS/MS counts show the first difference between workflows. A-UT runs acquired about half as many MS/MS spectra as the others (19,000–22,000 versus 40,000–51,000). The instrument therefore found fewer peptide precursors to select, even though A-UT gave the highest peptide concentration (Qubit). This suggests that deoxycholate and residual chaotrope, which A-UT does not remove before injection, suppressed ionisation or co-eluted with peptides. Acetone precipitation (A-PPT) and S-Trap washing (BST) remove both and doubled the number of spectra.

BST showed the most consistent performance across specimens, with 2,287 and 2,107 PSMs per run in MNRJ 24305 and MNRJ 24314, respectively. A-PPT performed similarly to BST in MNRJ 24305 but yielded about half as many PSMs in MNRJ 24314. This specimen-dependent performance may reflect differences in the amounts of matrix components and salts co-precipitated with the proteins during acetone precipitation.

BGEL generated the largest number of MS/MS spectra, but a substantial proportion of the assigned spectra came from contaminants. In MNRJ 24305, 57% of assigned spectra matched human keratins and trypsin introduced during sample preparation and gel handling, compared with 1–3% for the in-solution workflows; the corresponding proportion in MNRJ 24314 was 15%. After contaminant removal, BGEL gave intermediate depth, above A-UT but below A-PPT and BST (Table 1). Pooling the peptide extracts of all ten gel slices before injection may also favour repeated sampling of abundant proteins at the expense of distinct peptide sequences.

**Table 1.** Identification performance per run (mean of injections), LEMTEN pipeline, 1% FDR.

| Specimen | Workflow | Runs | MS/MS per run | PSMs per run | Assigned spectra per run | Contaminant spectra (%) | Protein groups per run |
| --- | --- | --- | --- | --- | --- | --- | --- |
| MNRJ 24305 | A-UT | 6 | 22,006 | 979 | 1,041 | 1.0 | 547 |
| MNRJ 24305 | A-PPT | 6 | 46,621 | 2,252 | 2,457 | 1.3 | 1,379 |
| MNRJ 24305 | BST | 6 | 43,927 | 2,287 | 2,401 | 2.6 | 1,222 |
| MNRJ 24305 | BGEL | 3 | 50,953 | 1,667 | 1,698 | 57.4 | 951 |
| MNRJ 24314 | A-UT | 6 | 18,717 | 533 | 564 | 0.7 | 369 |
| MNRJ 24314 | A-PPT | 6 | 39,934 | 1,091 | 1,178 | 2.6 | 730 |
| MNRJ 24314 | BST | 6 | 44,946 | 2,107 | 2,248 | 2.9 | 1,204 |
| MNRJ 24314 | BGEL | 3 | 49,453 | 1,183 | 1,240 | 15.4 | 684 |

PSMs (database search) and assigned spectra (adding homology and sequence-tag hits) exclude contaminant-only matches. The contaminant column gives their share of all assigned spectra.
Protein groups are inferred from the combined evidence, excluding contaminants.

**Table 2.** Same-database comparison, mean per run, 1% FDR, contaminants excluded. LEMTEN columns count only evidence explained by DB-1 sequences.

| Specimen | Workflow | FragPipe spectra | LEM TEN PSMs | LEM TEN PSMs + homology | FragPipe proteins | FragPipe proteins, $\geq 1$ unique spectrum | LEM TEN protein groups |
| --- | --- | --- | --- | --- | --- | --- | --- |
| MNRJ 24305 | A-UT | 1,220 | 856 | 911 | 523 | 212 | 468 |
| MNRJ 24305 | A-PPT | 2,769 | 1,969 | 2,146 | 1,296 | 560 | 1,202 |
| MNRJ 24305 | BST | 3,019 | 1,994 | 2,094 | 1,218 | 526 | 1,059 |
| MNRJ 24305 | BGEL | 1,967 | 1,411 | 1,438 | 876 | 364 | 787 |
| MNRJ 24314 | A-UT | 655 | 475 | 502 | 382 | 149 | 324 |
| MNRJ 24314 | A-PPT | 1,396 | 976 | 1,054 | 743 | 301 | 647 |
| MNRJ 24314 | BST | 2,753 | 1,850 | 1,968 | 1,218 | 522 | 1,054 |
| MNRJ 24314 | BGEL | 1,503 | 1,081 | 1,132 | 695 | 308 | 600 |

FragPipe spectra are protein spectral counts from combined\_protein.tsv.

Technical quality metrics were comparable across workflows. Median precursor mass errors ranged from 2.1 to 5.3 ppm, and 88–97% of identified peptides had no missed cleavage (median 95%). Across workflows, 75–81% of protein groups were supported by a single peptide, as is typical for metaproteomes searched against databases of distantly related sequences.

### S-Trap Gives the Deepest and Most Specimen-Independent Proteome

S-Trap (BST) combined high protein-group coverage with the smallest difference between specimens.

#### Protein groups per specimen

BST identified 2,196 protein groups in MNRJ 24305 and 2,025 in MNRJ 24314, an 8% difference. The other workflows lost 21–39% between specimens: A-PPT identified 2,330 and 1,421 groups, A-UT 988 and 627, and BGEL 1,091 and 866 in MNRJ 24305 and MNRJ 24314, respectively. These counts are unions over injections and extractions, requiring at least one spectrum and excluding contaminants.

#### Specimen means

Across specimen means, BST yielded 2,390 ± 105 assigned spectra and 1,213 ± 13 protein groups per run. A-PPT yielded 1,849 ± 905 assigned spectra, and BGEL 2,725 ± 1,782. The SAF rank–abundance curves showed the same pattern (Figure 3): the BST curves of the two specimens overlapped closely along their whole length, whereas the A-PPT and A-UT curves diverged after about the first 100 proteins.

**Figure 3.**
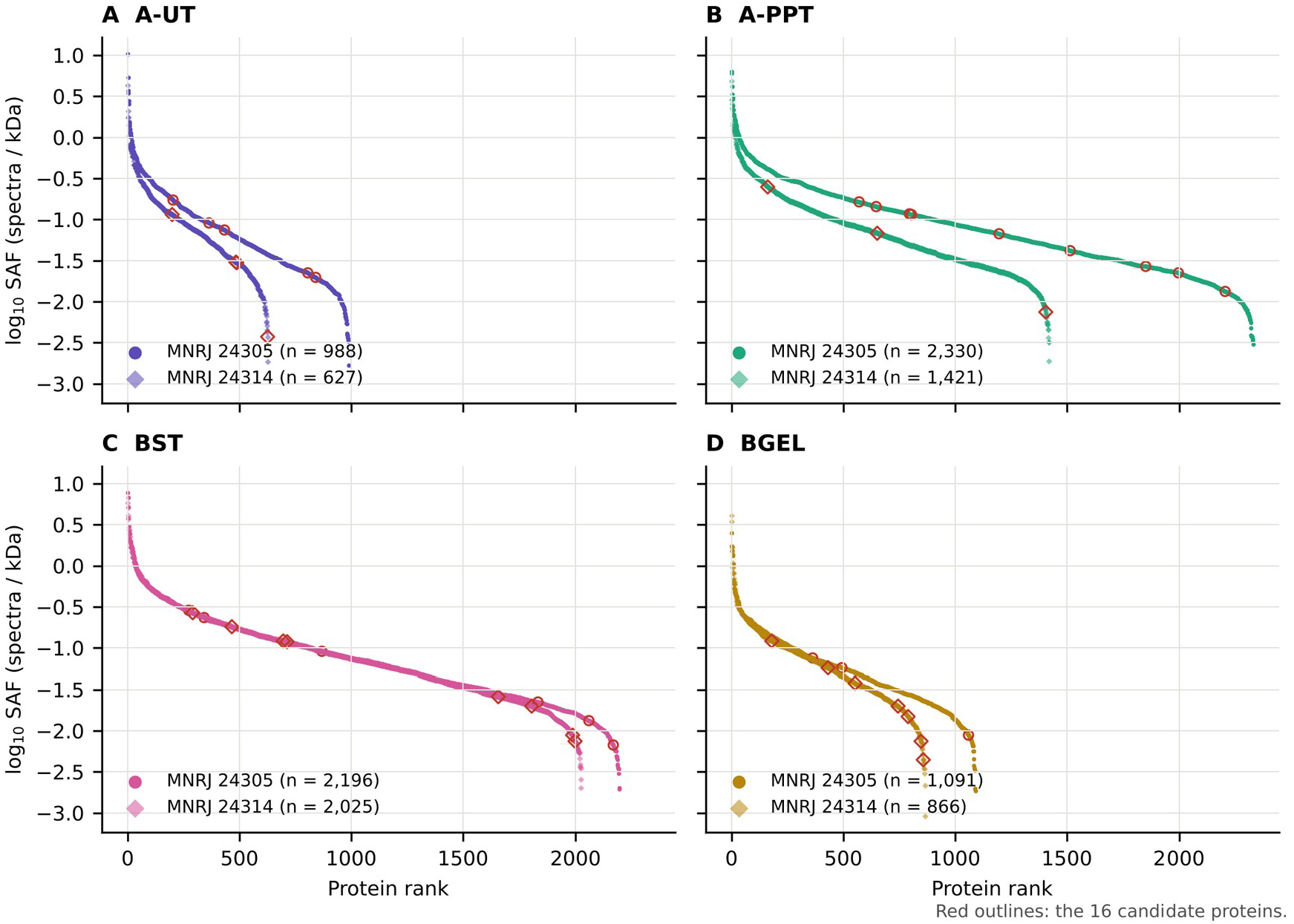
SAF rank–abundance curves of non-contaminant protein groups for (A) A-UT, (B) A-PPT, (C) BST and (D) BGEL, for MNRJ 24305 (circles) and MNRJ 24314 (diamonds). Curve length shows coverage. The two BST curves superimpose.

FragPipe gave the same within-specimen ranking with the stricter requirement of at least one unique spectrum. In MNRJ 24305, A-PPT identified 1,047 proteins, followed by BST (1,009), BGEL (498) and A-UT (438). In MNRJ 24314, BST identified 945 proteins, followed by A-PPT (674), BGEL (435) and A-UT (308). The original analysis reported 1,085, 1,064, 508 and 454 proteins for A-PPT, BST, BGEL and A-UT, respectively, in MNRJ 24305, and 704, 954, 435 and 317 in MNRJ 24314.

#### Overlap between workflows

The harmonised comparison included 4,841 protein groups, of which 2,087 were detected in both specimens by at least one workflow and 254 by all four workflows (Figure 4A). BST contributed the largest workflow-exclusive set (509 groups), followed by A-PPT (279), BGEL (65) and A-UT (41). A further 410 groups were shared only by A-PPT and BST. The two workflows that remove detergent and chaotrope before digestion therefore recover a common set of proteins that is less represented in the other workflows, and BST adds the largest set of workflow-specific identifications.

**Figure 4.**
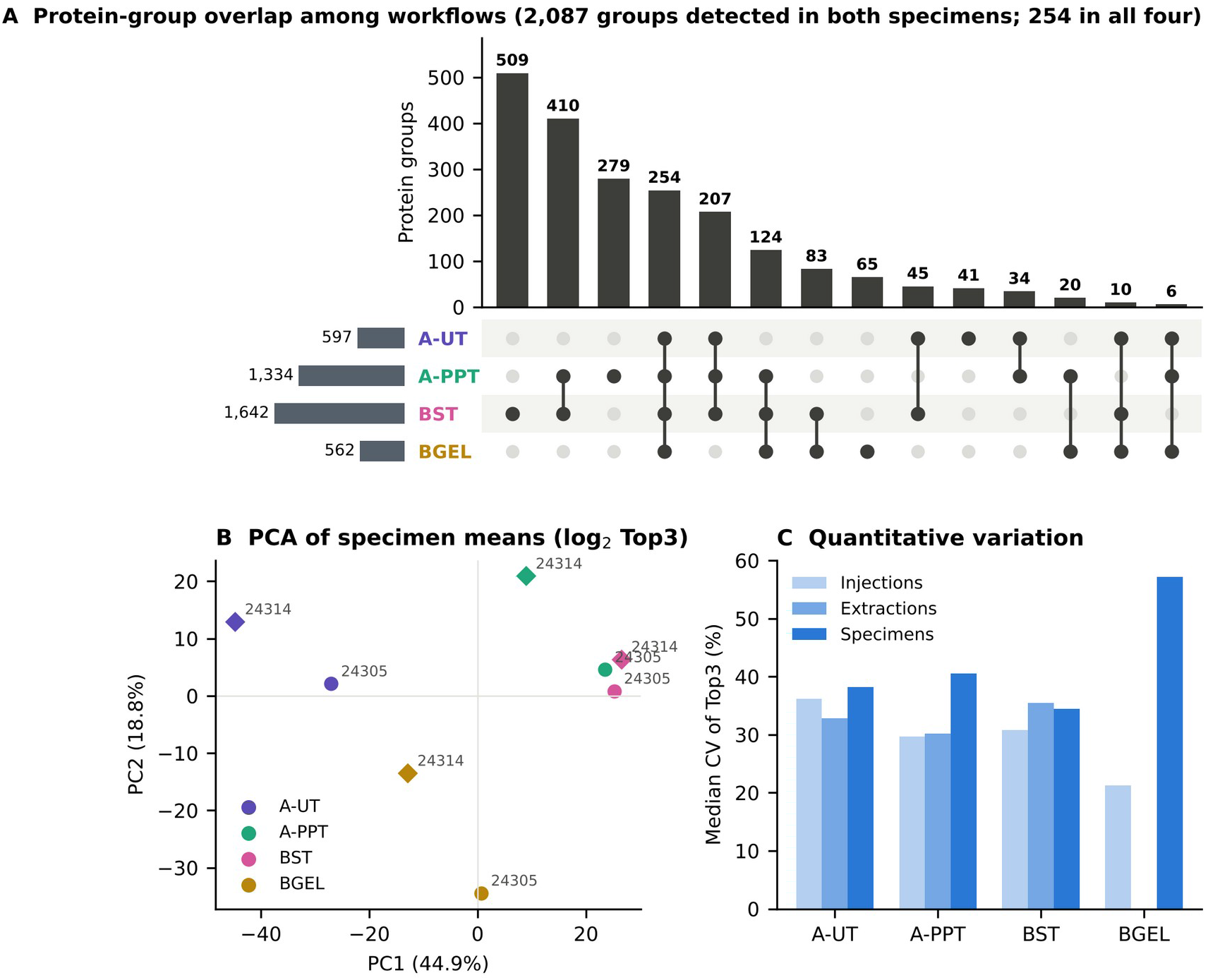
Overlap, structure and reproducibility. (A) UpSet plot of protein groups shared among workflows. A group counts for a workflow when it was detected in both specimens; 2,087 of the 4,841 groups met this in at least one workflow, and 254 in all four. Top bars, groups in each combination (filled dots); left bars, groups per workflow. (B) PCA of log2 Top3 abundance averaged per specimen. (C) Median coefficients of variation between injections, extractions and specimens per workflow. The 47 groups differing between workflows (ANOVA, q < 0.05) are shown as a heatmap in Figure S4.

A-UT contributed little additional coverage. Its 41 exclusive groups were fewer than the 65 of BGEL, although A-UT was analysed in twice as many runs.

#### Quantitative reproducibility

Median coefficients of variation (CVs) of Top3 abundance ranged from 30% to 36% between extractions and from 21% to 36% between injections (Figure 4C). BGEL showed the lowest between-injection CV (21.3%), as its three injections came from a single pooled digest. Among the in-solution workflows, A-PPT and BST had the lowest between-injection CVs (29.7% and 30.8%), and A-UT the highest (36.2%).

BST also had the lowest between-specimen CV (34.4%), compared with 38.2% for A-UT, 40.5% for A-PPT and 57.2% for BGEL. Correlations between duplicate extractions were r = 0.82–0.83 for BST, 0.77–0.81 for A-PPT and 0.77–0.79 for A-UT. These values are lower than the MaxLFQ correlations obtained with FragPipe (r = 0.90–0.97); the two rest on different quantification approaches, and Top3 abundances from single injections are expected to vary more.

#### Principal components

In the PCA of specimen means (Figure 4B), PC1 (44.9% of the variance) separated A-UT from A-PPT and BST, and PC2 (18.8%) distinguished BGEL from the other workflows. In the run-level PCA, PC1 explained 40.8% of the variance, and injections of the same preparation clustered closely. The 12 BST runs of both specimens formed a single cluster, whereas A-PPT runs separated by specimen.

Among the 254 protein groups common to all workflows, 47 differed significantly between workflows (ANOVA on specimen means, q < 0.05). They followed three main patterns. The first comprised proteins less abundant with A-UT, including ADP-ribosylation factor 6, α-actinin, annexin A4 and elongation factor Tu, which may reflect incomplete solubilisation or interference from residual detergent. The second comprised proteins more abundant with BGEL, including the cell-surface protein SprA, TonB-dependent receptors and periplasmic transporters assigned to Poribacteria; the gel-based workflow may favour the recovery of large, membrane-associated bacterial proteins that in-solution digestion handles less efficiently. The third comprised specimen-dependent patterns of EF-hand and IPT/TIG domain proteins and the 60 kDa chaperonin, which were less abundant with BGEL. BGEL may therefore complement BST for selected symbiont envelope proteins rather than replace it.

#### Candidate proteins

The 16 candidate proteins selected in our preliminary analysis of these data were evaluated in both pipelines, and those detected by both generally showed consistent workflow patterns (Table 3). Talin-1 (A0AA35WUK6) and the TIR-domain protein (A0A1X7V981) were detected in all eight specimen × workflow combinations by both pipelines, and BST gave the highest spectral counts for talin-1 in both specimens. Two candidates, the integrin β homologue O18482 and A0AA96S0V6, had no assigned spectra in either pipeline; differences in accession formats between database versions may contribute to this absence.

**Table 3.**
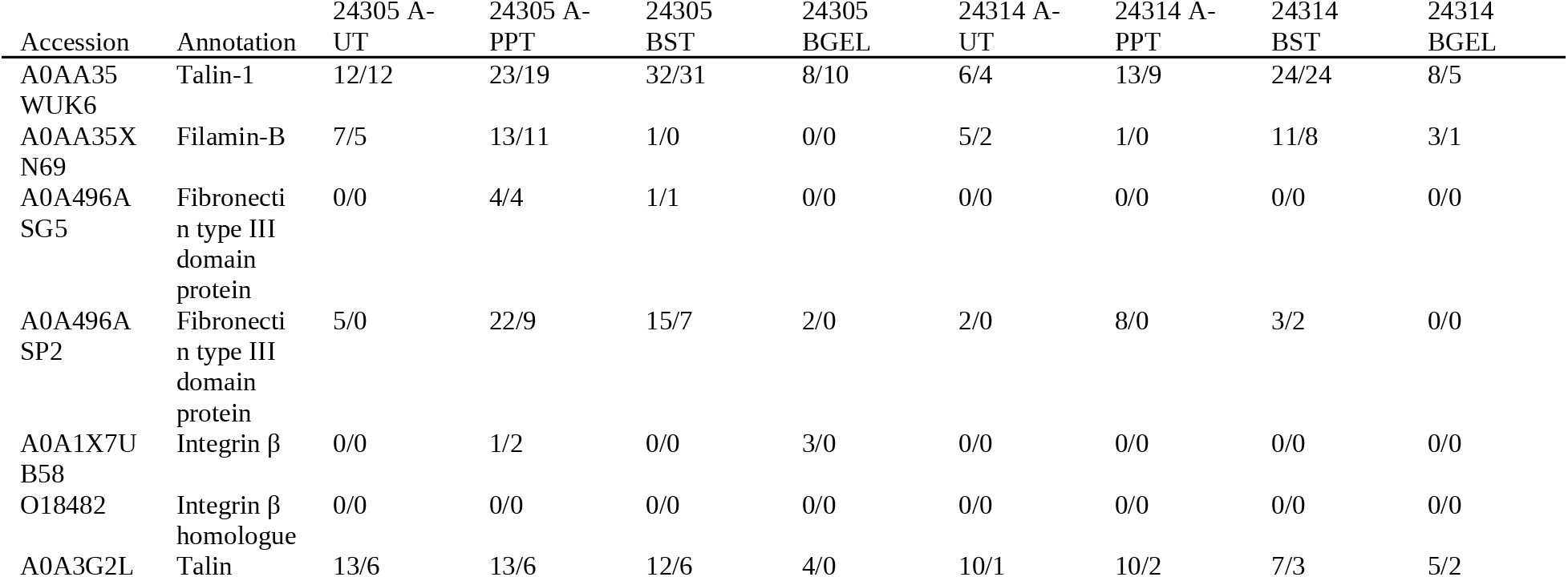
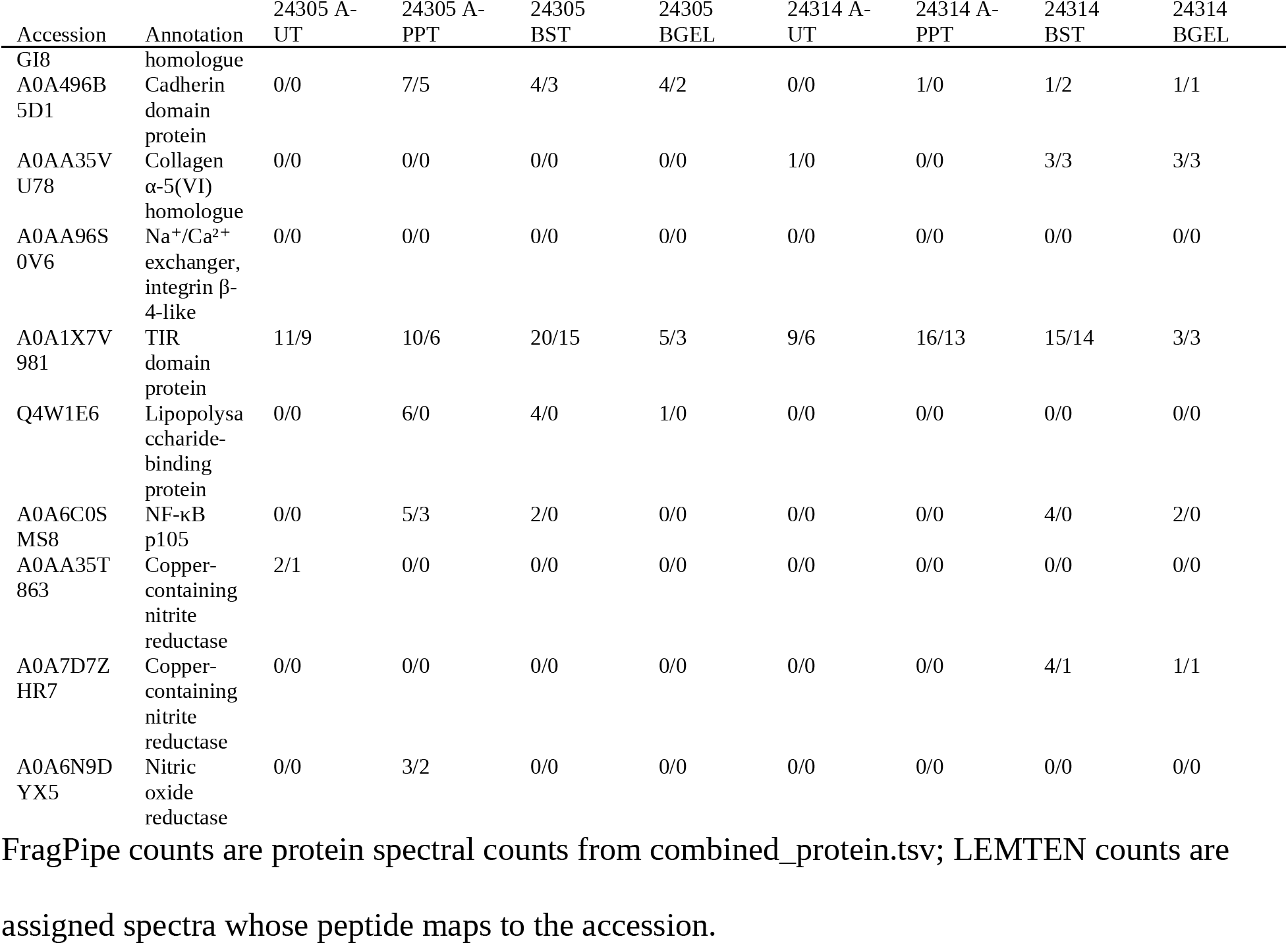
Spectral support for the 16 candidate proteins, FragPipe (DB-1) / LEMTEN (DB-2) spectra summed over the runs of each specimen × workflow.

LEMTEN counts were generally equal to or lower than FragPipe counts, which is expected because *H. panicea* homologues absorb shared peptides in DB-2. The lipopolysaccharide-binding protein Q4W1E6 was supported only by FragPipe, consistent with the reassignment of its peptides to an *H. panicea* homologue in LEMTEN.

#### Workflow selection for phase 2

S-Trap combined consistently high identification depth, the smallest difference between specimens and the largest number of workflow-exclusive protein groups. It was therefore carried forward to the biological quadruplicate of phase 2.

### On the Same Database, FragPipe Is Deeper and the Two Pipelines Agree

The two pipelines were compared on equal terms. FragPipe searched DB-1, whereas the LEMTEN pipeline searched DB-2, which adds the *H. panicea* proteome. LEMTEN identifications were therefore restricted to those explained by DB-1 sequences: spectra whose peptides occur only in *H. panicea* proteins were removed, and contaminant-only matches were excluded from both outputs.

As a direct check of this restriction, Sage was also re-run on DB-1 alone for six runs covering all workflows and both specimens (Supporting Table S5). The re-search gave 2–10% more PSMs than the exclusion estimate (median 7%; 625–1,977 PSMs per run), so the exclusion is slightly conservative and does not favour the LEMTEN pipeline. On these six runs, Sage PSMs on DB-1 were 64–77% of the FragPipe spectral counts.

On equal sequences, the two strategies ranked the 42 runs identically (Spearman ρ = 0.98 between per-run FragPipe spectral counts and LEMTEN DB-1 PSMs). The workflow ranking is therefore not an artefact of the search engine.

At the PSM level, FragPipe gave greater identification depth. With Sage alone, LEMTEN recovered a median of 70% of the FragPipe spectral counts per run, rising to 75% after adding homology hits on DB-1 sequences. Because summed protein spectral counts overstate the number of distinct PSMs when peptides are shared between proteins, two BST runs of MNRJ 24305 (extraction 1, injections 2 and 3) were also compared scan by scan. FragPipe reported 2,843 and 2,590 PSMs in these runs, against 1,869 and 1,790 Sage PSMs from the DB-1 re-search. Of the 1,688 and 1,586 scans identified by both, 98.7% and 99.2% received the same peptide, treating isoleucine and leucine as equivalent. FragPipe alone identified 1,155 and 1,004 scans, and Sage alone 181 and 204. On the same sequences, MSFragger with MSBooster predictions and Percolator rescoring therefore identified about 1.4–1.5 times as many spectra as Sage with our SVM rescoring, while the assignments of scans identified by both were highly concordant.

At the protein level the totals were closer: LEMTEN protein groups per run were a median of 89% of all FragPipe proteins. Requiring at least one unique spectrum, as in the original analysis, retained about 40% of the FragPipe proteins. This rule excludes proteins supported only by shared peptides, including members of the large Porifera protein families in the database. The main contribution of the LEMTEN pipeline therefore lies in extending the evidence obtainable from a fixed-database search, not in conventional sensitivity. This additional evidence comes from the proteome of the closest sequenced relative, from sequence-divergent peptides and from semi-specific peptides (next section). For non-model organisms, these results support combining the most sensitive conventional search available with homology-driven layers.

### Homology-Driven Search Adds One-Fifth More Evidence

The homology-driven layers of the pipeline increased the number of assigned non-contaminant spectra by 21.3% relative to a conventional search of the laboratory database. A Sage search of DB-1 sequences identified 56,199 spectra across the 42 runs, and three further layers brought the total to 68,150 (Figure 5). Adding the *H. panicea* proteome yielded 7,842 PSMs whose peptides occur only in its proteins (+14.0%), covering 812 distinct peptides absent from every other database entry. Two homology layers extended these identifications: alignment of de novo sequence tags to DB-1 with PAM30MS, followed by precursor-mass rescoring, added 3,559 spectra, and the same procedure against *H. panicea* proteins added 550.

**Figure 5.**
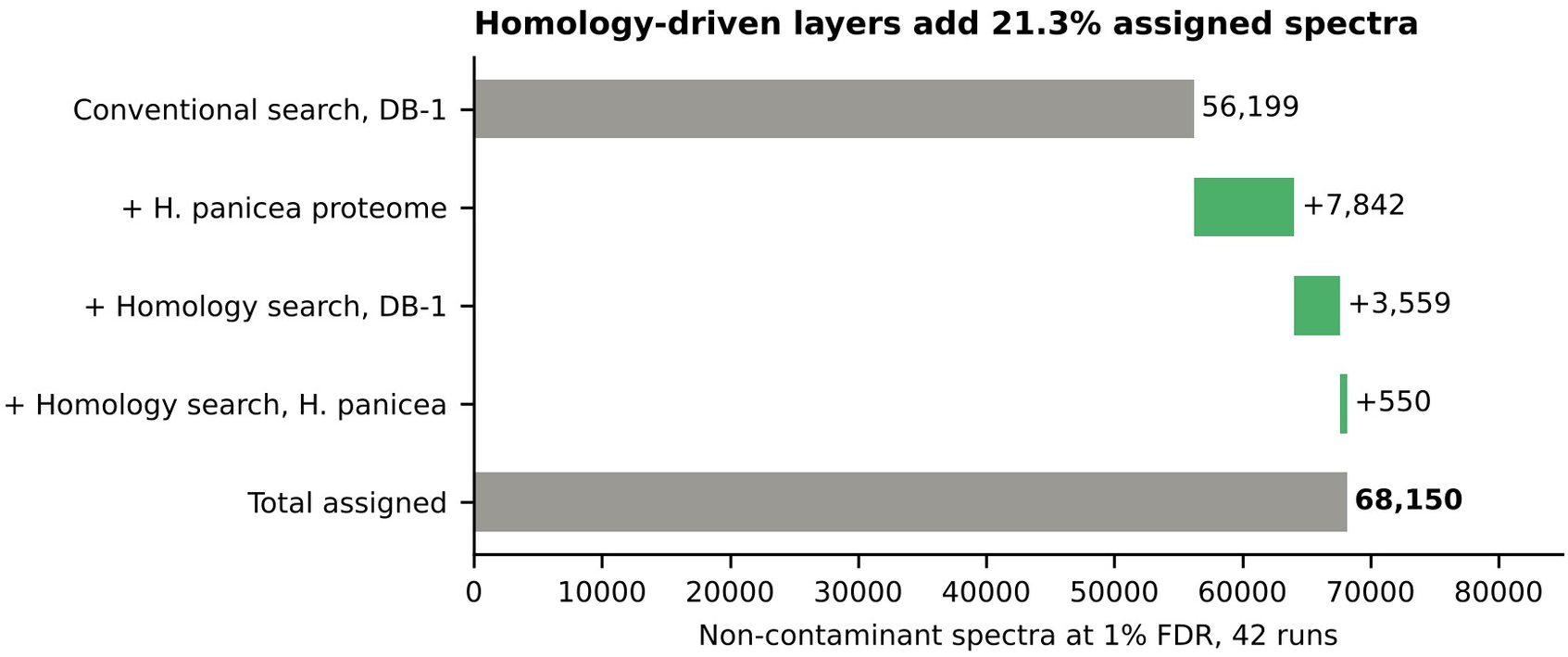
Spectra gained by each homology-driven layer over a conventional search of DB-1, summed over 42 runs at 1% FDR.

*H. panicea* proteins became the leading proteins of 14.6% of all protein groups. A single genome from a related family thus contributed about one in nine assigned spectra, on top of a database of about 474,000 sequences. That database is dominated by *Geodia barretti* and *Amphimedon queenslandica*, which belong to more distantly related orders, so a phylogenetically closer reference proteome added evidence that a much larger but more distant collection could not. The agreement between homology-based and conventional assignments was evaluated on spectra both approaches could explain. Across the 42 runs, with the FDR estimated over all spectra, the homology search assigned 23,684 spectra at 1% FDR, of which 16,626 also had a Sage PSM. For 99.5% of these shared spectra (95.7–100% per run), both approaches reported the same peptide, treating isoleucine and leucine as equivalent.

The other 7,058 had no Sage PSM. For the combined result, the homology search was repeated on spectra without a Sage PSM only, with its own target–decoy estimate. Because this subset holds lower-quality spectra, the threshold is stricter: 4,323 spectra passed 1% FDR, 214 of them matching only contaminants, and the remaining 4,109 non-contaminant spectra form the two homology layers above (3,559 + 550). Of the 4,323, 2,643 matched a database peptide exactly but scored below the 1% FDR threshold in the Sage search; the de novo tag and precursor mass recovered them. The remaining 1,680 were homologous peptides that differ from every database sequence by amino-acid substitutions or a mass shift, which a conventional search cannot report. These homology spectra added 3,740 peptides to the per-run peptide lists and supported 1,969 protein groups identified only by homology (5.2% of all groups): 1,514 sponge and 455 symbiont groups. The relative gain ranged from 2.1% to 9.0% across specimens and workflows. It was highest for A-PPT (+9.0% in MNRJ 24305 and +7.9% in MNRJ 24314), intermediate for A-UT and BST (+4.9% to +6.5%) and lowest for BGEL (+2.1% and +4.6%). The gain therefore did not simply follow the depth of the conventional search.

The error-tolerant sequence tag identified a further class of spectra. Among hits at q ≤ 0.05, the unaligned flank was resolved as a semi-specific peptide in 436 target hits, against 5 decoy hits (decoy-to-target ratio 1.1%). Such assignments may reflect non-tryptic cleavage or in-source fragmentation, and they lie outside the fully tryptic specificity of the conventional search.

More broadly, 4,571 target hits had a precursor mass difference not explained by the matched database peptide, against 47 decoy hits (decoy-to-target ratio 1.0%). The most frequent shifts recurred in most runs and were almost absent among decoys: −1.97 Da (T→V), 367 target versus no decoy hits, in 27 runs; −14.01 Da (A→G, T→S, L→V), 225 versus 1, in 35 runs; +13.02 Da (D→Q), 165 versus 1; +15.00 Da (N→E), 115 versus 0; +0.98 Da (deamidation, N→D or Q→E), 78 versus 0; and −15.99 Da (S→A, Y→F), 36 versus 0 (Figure 6). Nearly all correspond to single amino-acid substitutions between *V. ophiraphidites* and its database homologues, which shows, spectrum by spectrum, the sequence divergence that limits a conventional search. In addition, Unimod modifications were localised to an unaligned flank in 101 target hits.

**Figure 6.**
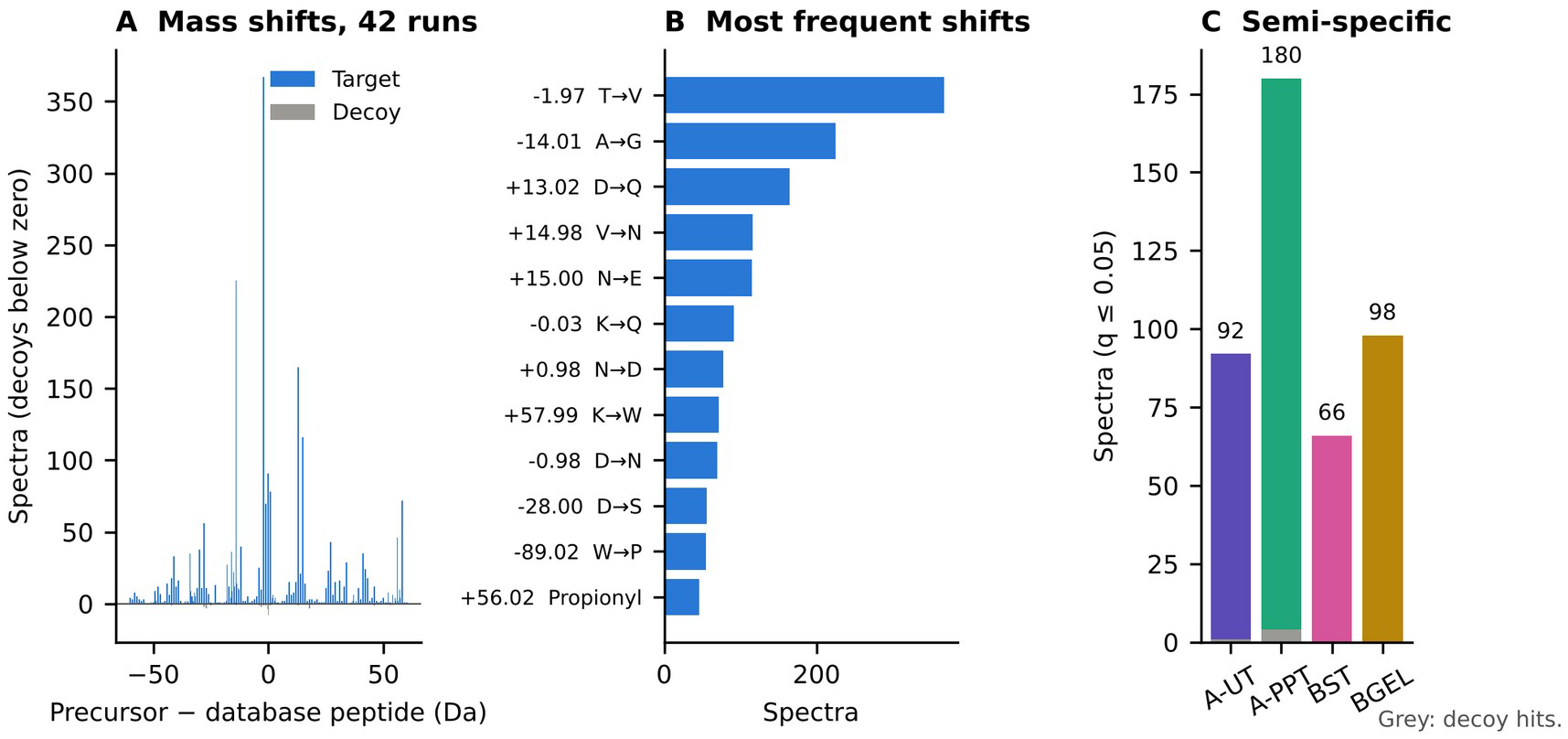
Mass-shift panorama of homology hits (q ≤ 0.05). (A) Distribution of precursor–database-peptide mass differences for targets and decoys. (B) Most frequent shifts with their annotation (single amino-acid substitutions, Unimod modifications, isotope errors). (C) Semi-specific peptides explained by the error-tolerant sequence tag, with target and decoy counts.

The FDR of the homology layer was checked against a positive control: spectra already identified by Sage were searched by homology as if unknown, and the share whose best homology hit disagreed with the Sage peptide gives the observed error. At a nominal 1% FDR, the observed error was 1.0% pooled over the 42 runs (24,759 control spectra; median 1.2% per run, range 0–4.0%). With the earlier version of the rescoring step, which reconstructed the decoys incorrectly (see Database Construction, Optimization and Testing), the same check gave 2.0%, and the homology layer reported 6,236 rather than 4,323 new spectra.

The contribution of the successive rescoring steps was evaluated on one BST run (MNRJ 24305, extraction 1, injection 1) by counting new spectra at 1% FDR: 141 with the PAM30MS alignment alone, 118 after precursor-mass rescoring, 106 after multivariate rescoring and 102 with the error-tolerant tag. With correct decoys, rescoring did not add spectra in this run. It raises decoy scores as much as target scores, so its role is to rank alignments by plausibility and to explain mass shifts, not to increase the count.

### A First Holobiont Proteome of *V. ophiraphidites*

Across all workflows and both specimens, the combined evidence identified 4,294 distinct protein-group leaders after excluding contaminants. Of these, 1,787 (42%) were assigned to sponge proteins and 2,507 (58%) to bacterial or archaeal proteins. These counts treat as bacterial the 717 leaders that DB-2 labels *Geodia barretti* but whose sequences are bacterial (see Database Construction, Optimization and Testing); with the database labels, the split would have been 58% sponge and 42% microbial. A set of 53 groups, 36 sponge and 17 symbiont proteins, was detected in at least 40 of the 42 runs of both specimens. These groups form an operational core proteome across the specimens and workflows examined, not a species-wide core proteome.

By spectra, the host still contributed most of the evidence: 33,931 of the 61,487 spectra assigned to non-contaminant leaders (55%), 29,457 of them to Demospongiae. Bacterial evidence was assigned predominantly to Poribacteria, a candidate phylum characteristic of high-microbial-abundance sponges^32^, which contributed 15,191 spectra and 1,272 groups. Leaders reclassified from the *Geodia* proteome added 8,728 spectra of unclassified Bacteria, led by 60 kDa chaperonins (GroEL) and elongation factor Tu. Nitrospirota (1,257 spectra) and Planctomycetota (973 spectra) followed, with smaller contributions from Pseudomonadota and Actinomycetota, and archaeal evidence was limited to a single protein assigned to Thermoproteota (Figure 7).

**Figure 7.**
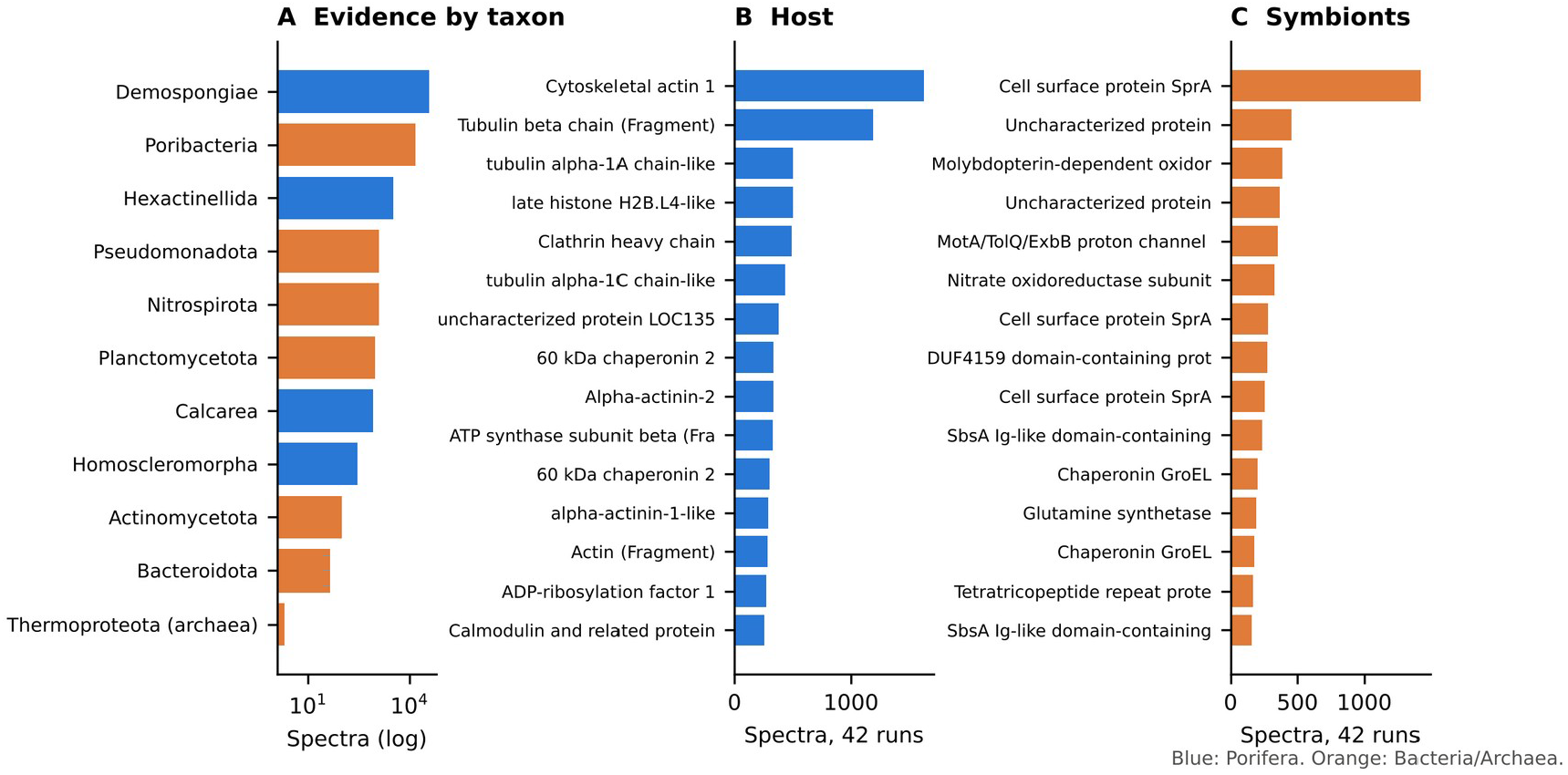
Taxonomic and functional composition of the *V. ophiraphidites* holobiont proteome. (A) Spectral evidence by taxon (log scale). Bacteria (unclassified): leaders labelled *Geodia barretti* in DB-2 whose sequences are bacterial. (B) The 15 most abundant host proteins (spectra, 42 runs). (C) The 15 most abundant symbiont proteins.

Leaders from Hexactinellida (3,345 spectra), Calcarea and Homoscleromorpha should not be read as evidence that these sponge classes were present. These entries are the best-scoring group members for highly conserved proteins, such as actin, histone H4 and ATP synthase, whose identical peptides occur across Porifera. Taxonomic interpretation below the class level therefore requires peptides that distinguish the relevant lineages, which are available only for divergent proteins.

#### Host

The most abundant host proteins were associated with the cytoskeleton, biosynthesis, protein homeostasis and energy metabolism. Cytoskeletal and trafficking proteins included actin, α- and β-tubulin, α-actinin, myosin heavy chains and clathrin heavy chain. Other abundant proteins were histones H2A, H2B and H4, HSP90, and ATP synthase α and β subunits. These identifications describe a host proteome rich in structural and housekeeping proteins, although abundance alone does not establish rates of cellular activity. Further host proteins included the pentose-phosphate enzymes transketolase and 6-phosphogluconate dehydrogenase, a 14-3-3 protein, and an H₂O₂-forming NAD(P)H oxidase possibly involved in reactive-oxygen signalling or defence.

Structural and host–microbe interaction candidates of the original analysis recovered by both pipelines included talin-1, a talin homologue, filamin-B, fibronectin type III domain proteins and the TIR-domain protein (Table 3). *H. panicea* proteins led 14.6% of groups, including many core host proteins that the UniProt sponge entries represent only as fragments.

#### Symbionts

The symbiont proteome contained prominent envelope proteins alongside proteins of nitrogen and carbon metabolism. Among proteins assigned to Poribacteria, the leading identifications were the cell-surface protein SprA, part of the Bacteroidetes-type secretion and gliding machinery, MotA/TolQ/ExbB channels, the outer-membrane lipoprotein BamD and SbsA Ig-like surface-layer proteins. These point to cell-surface and envelope functions that may contribute to interactions with the host, although they do not directly demonstrate attachment to the host matrix.

Proteins assigned to *Nitrospira* included the β subunit of nitrite oxidoreductase (NxrB), detected in all 42 runs, and a molybdopterin-dependent oxidoreductase, detected in 41. Their consistent detection provides protein-level evidence for nitrite-oxidation machinery in the holobiont. Poribacterial glutamine synthetase and a P-II nitrogen regulator further indicate nitrogen assimilation. Xanthine dehydrogenase family proteins suggest a contribution of purine catabolism, and phosphoenolpyruvate carboxykinase points to gluconeogenesis. Together, these proteins are consistent with a metabolically diverse symbiont community involved in nitrogen cycling, as described for other high-microbial-abundance sponges^4^. They show expressed components of these pathways, not pathway flux or nitrification rates.

#### Effect of workflow on the holobiont profile

Workflows influenced the apparent host–symbiont balance as well as the depth. The share of protein groups assigned to symbionts per run ranged, on average per workflow and specimen, from 39% (A-UT, MNRJ 24305) to 52% (BGEL, MNRJ 24314). Homology search added three times as many sponge groups as symbiont groups (1,514 versus 455). Sequence divergence between *V. ophiraphidites* and the database sponges was therefore the main limitation of the conventional search, with a larger impact on host than on symbiont identifications.

#### Peptide-level taxonomy

Because protein-level assignment depends on which database entry leads each group, taxonomy was also assessed at the peptide level. Each of the 8,086 distinct peptides was assigned to the lowest common ancestor (LCA) of all database proteins containing its sequence (Supporting Figure S6). Of these, 3,544 (43.8%) were assigned to Porifera, 3,106 of them resolving to Demospongiae, and 4,480 (55.4%) to Bacteria. Within the bacterial fraction, Poribacteria held 2,175 peptides (49%), and 1,532 could not be placed below the domain, almost all of them from bacterial entries of the *Geodia* proteome. Planctomycetota (345), Pseudomonadota (266) and Nitrospirota (127, all Nitrospiria) followed. Archaeal assignments comprised 13 peptides, 12 of them from Nitrososphaeria, the class of ammonia-oxidizing archaea. Together with *Nitrospira* NxrB, these assignments are consistent with both steps of nitrification in the holobiont, although taxonomic assignments alone do not establish their activity. The remaining 49 peptides (0.6%) were shared across domains and assigned to the root. The peptide-level view agrees with the protein-level picture once the bacterial *Geodia* entries are relabelled: the symbionts contribute about as many distinct peptides and protein groups as the host, and Poribacteria dominate the assignable bacterial fraction. The host contributes more spectra (55%), because its most abundant proteins (actin, tubulin, histones) are sampled many times. The 167 peptides resolving to Hexactinellida require the same caution as the protein-group leaders: they reflect the database sponge that holds the matching sequence, not the presence of glass sponges.

### Phase 2: Four Specimens Support a Shared Holobiont Signature and Reveal Preservation-Associated Differences

In phase 2, S-Trap was applied to four specimens: MNRJ 24305 and MNRJ 24314, analysed in phase 1, and two additional specimens, MNRJ 24317 and MNRJ 24325 (Figure 8). This analysis tested whether the workflow performance and holobiont profile of phase 1 held across individuals. The four specimens also differed in preservation history, which allowed an exploratory comparison of frozen and ethanol-stored material, although preservation was confounded with collection site and storage history.

**Figure 8.**
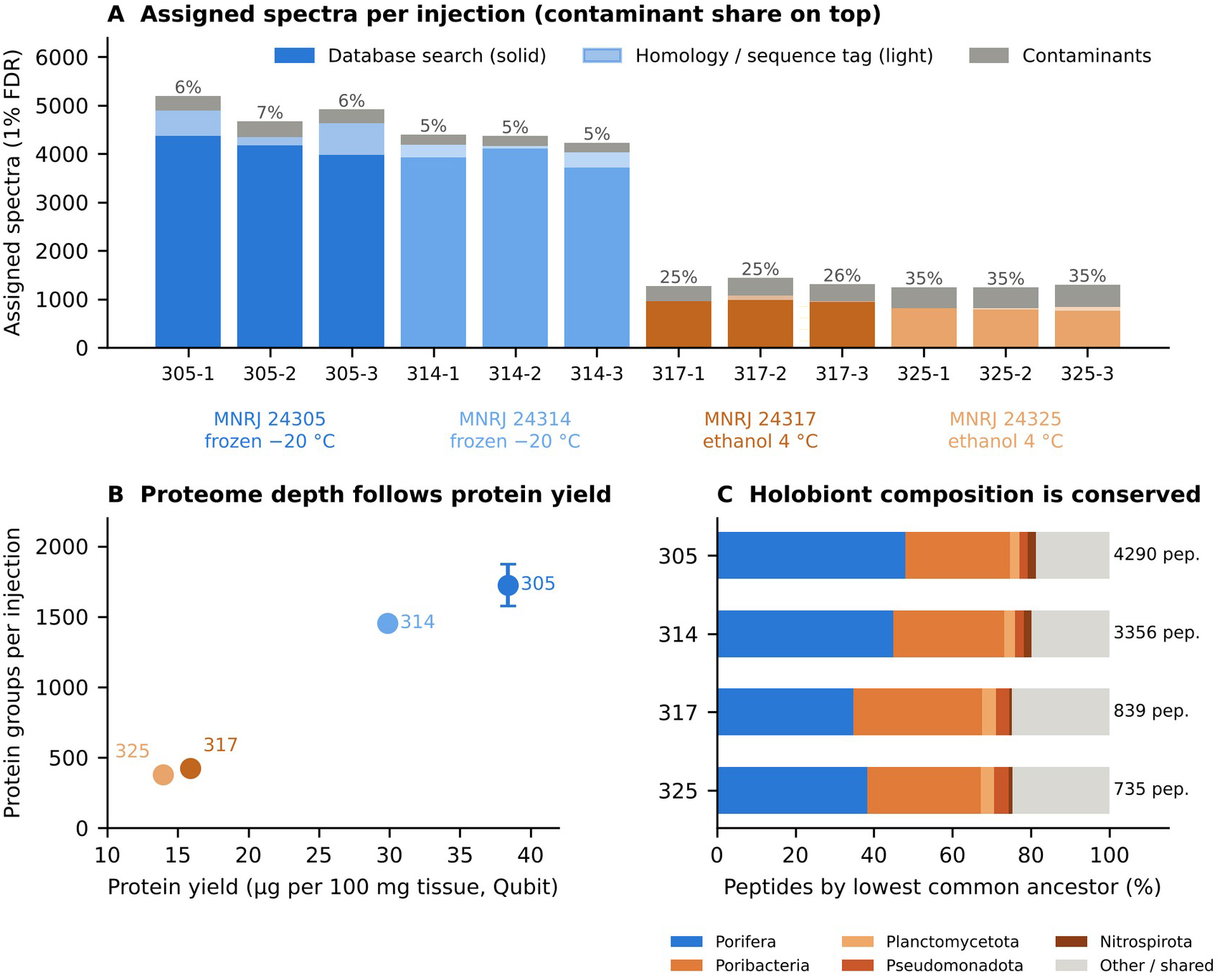
Biological quadruplicate: four specimens of *V. ophiraphidites*, each digested by S-Trap and injected in technical triplicate. (A) Assigned spectra per injection at 1% FDR. Bar colour identifies the specimen: blues, frozen at −20 °C (MNRJ 24305, dark; MNRJ 24314, light); oranges, ethanol at 4 °C (MNRJ 24317, dark; MNRJ 24325, light). Within each bar, the solid segment is database-search PSMs, the light segment homology or sequence-tag spectra, and the grey segment contaminants (keratins, trypsin), whose share is shown above the bar. (B) Protein groups per injection (mean ± SD) against protein yield per 100 mg of tissue, in the same specimen colours. (C) Taxonomic composition by peptide lowest common ancestor for each specimen: Porifera, blue; Poribacteria, Planctomycetota, Pseudomonadota and Nitrospirota, shades of orange to brown; other or cross-domain peptides, light grey.

#### Reproducibility and depth

Injections of the same digest were reproducible in all four specimens, with median spectral-count CVs of 22–25% (Supporting Figure S5D). In the PCA of the 12 runs, injections clustered by specimen, and PC1, which explained 70% of the variance, separated frozen from ethanol-stored specimens (Supporting Figure S5C). Identification depth, however, differed four- to fivefold between specimens. The specimens stored frozen at −20 °C, MNRJ 24305 and MNRJ 24314, yielded 4,036–4,889 assigned non-contaminant spectra and 1,412–1,896 protein groups per injection, with contaminants accounting for 5–7% of assigned spectra. The specimens stored in ethanol at 4 °C, MNRJ 24317 and MNRJ 24325, yielded 810–1,077 assigned spectra and 374–455 groups, with contaminants accounting for 25–36% (Supporting Table S6). Consistent reinjection performance therefore did not remove the differences in recoverable evidence between specimens.

#### What limits the ethanol-stored specimens

All 12 injections acquired similar numbers of MS/MS spectra (50,997–62,587), so the lower yield of the ethanol-stored specimens was not due to fewer acquired spectra. The keratin and trypsin background was similar or larger in these specimens (318–461 versus 194–323 spectra), whereas sponge and symbiont assignments fell four- to fivefold. Protein recovery was also lower: 15.9 and 14.0 µg per 100 mg of tissue, against 38.4 and 29.9 µg for the frozen specimens. Equal peptide loading by Qubit therefore did not translate into comparable amounts of sample-derived peptide, and identification depth tracked protein recovery (Figure 8B).

Preservation method, collection site and storage history were confounded in this set: both ethanol-stored specimens came from Sapata, were collected on the same day and were kept in the museum collection rather than in our laboratory. The lower coverage cannot therefore be attributed specifically to ethanol preservation. Nevertheless, the frozen specimens gave substantially more evidence under the conditions examined, which argues for frozen storage of sponge tissue at −20 °C rather than long-term storage in ethanol at 4 °C, and for checking protein yield and the contaminant share before comparing specimens.

#### Alkylation failed in the two frozen specimens

The problem was first noticed in an exploratory reanalysis by a naive LLM agent session that searched MNRJ 24305 without fixed carbamidomethylation, found more identifications than the standard search and almost no cysteine-containing peptides, and asked whether the samples had been alkylated^27^. A search of all 54 runs with the cysteine forms as variable modifications answered the question (Supporting Table S9). In the two frozen phase-2 specimens (MNRJ 24305 and MNRJ 24314), cysteine occurred in 0.6–1.0% of PSMs, against 3–10% in phase-1 S-Trap injections of the same specimens; most cysteines were disulfide-bonded (48–64%) or free (14–21%), and only 3–12% were carbamidomethylated. These two preparations were therefore not reduced and alkylated, for reasons that could not be established. The two ethanol-stored specimens of the same phase were alkylated (77–98% of cysteines carbamidomethylated). Because the two unalkylated preparations are exactly the two specimens with a different history, kept frozen in our laboratory before the equalisation step, a cause linked to that history cannot be excluded. In phase 1, 63–98% of cysteines were carbamidomethylated in S-Trap and acetone-precipitation preparations, with incomplete reduction (9–22% disulfide-bonded) in three of the fourteen preparations (MNRJ 24305 BST extraction 2, and A-PPT extraction 1 of both specimens). GeLC preparations also carried propionamide cysteine from the gel, as expected, and urea/thiourea preparations yielded few cysteine peptides (0.1–2.4% of PSMs), most of them carbamidomethylated.

Because the searches used fixed carbamidomethylation, the counts of the two frozen phase-2 specimens lack their cysteine peptides. The loss is small next to the difference between storage groups, since the frozen specimens gave four to five times the evidence despite it. Storage is nonetheless also confounded with sample preparation in phase 2.

#### Search performance

The searches behaved as in the 42-run phase-1 set. The homology-driven layers increased assigned spectra by 26.8%, from 24,992 to 31,702 (Supporting Figure S5A). Most of this gain came from the *H. panicea* proteome (4,529 spectra), and the homology layers added 2,181; their contribution varied widely between injections of the same digest (0–651 spectra), as in phase 1. Where homology and database searches explained the same spectrum, they agreed on 99.9% of peptides. The error-tolerant tag added 179 semi-specific peptides, with no decoy hits, and FragPipe on the same database ranked the runs in the same order (Spearman ρ = 0.97).

#### A shared holobiont signature

Despite the differences in depth, the peptide-level taxonomic composition was similar across specimens (Figures 8C and 9; Supporting Table S8). Bacteria accounted for 51–64% of peptides, with Poribacteria alone contributing 27–33%; a further 18–24% could be placed only at the domain level, including the peptides of the *Geodia* entries relabelled as bacterial. Porifera accounted for 35–48% of peptides, less in the ethanol-stored specimens. Nitrospirota, Planctomycetota and Pseudomonadota were detected in all four specimens, each at 1–4% of peptides, and Archaea at 0.1–0.4%. These shared proportions support a broadly conserved host–symbiont profile among the individuals examined, although they do not establish identical community composition or exclude differences in less abundant taxa.

**Figure 9.**
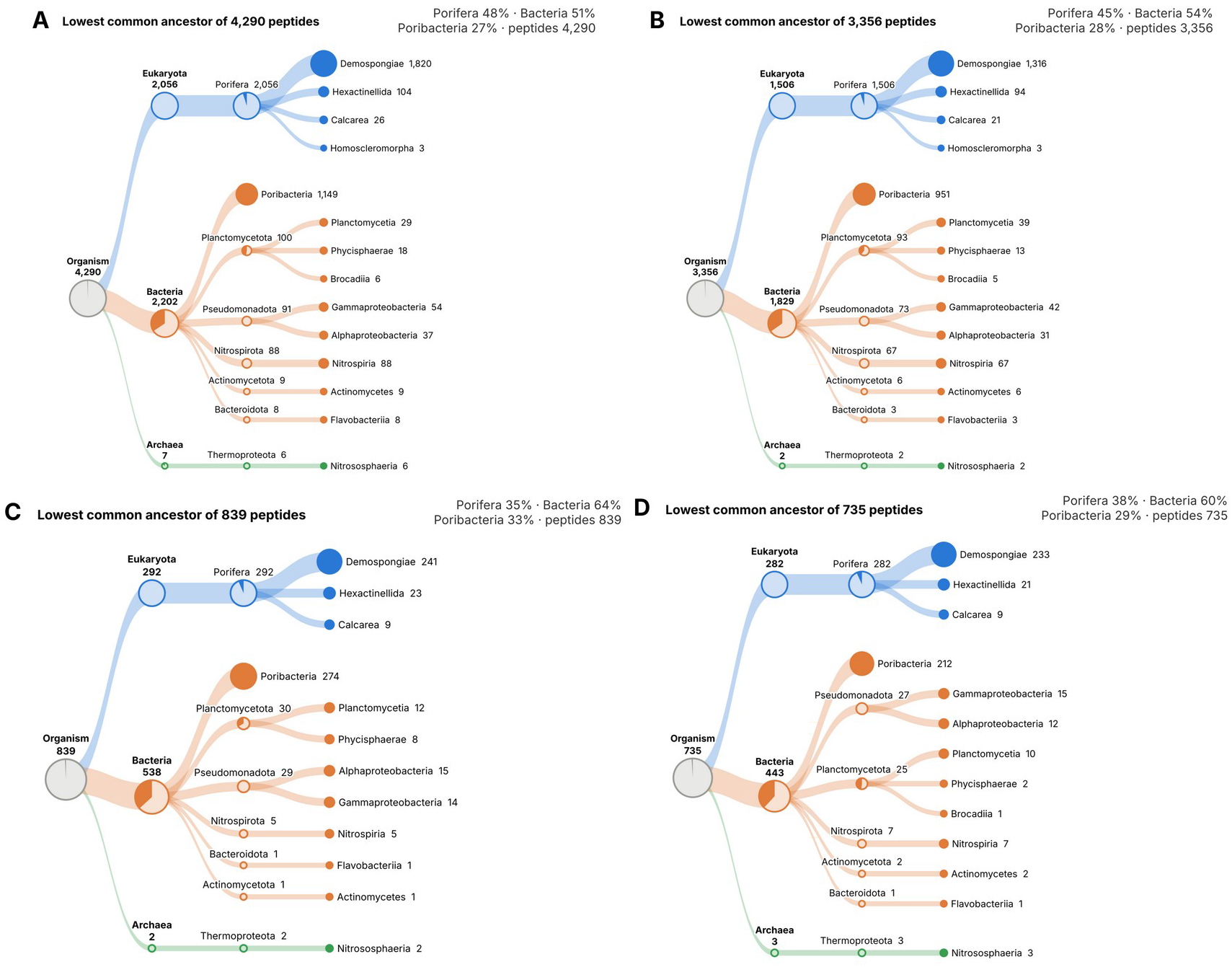
Taxonomic origin of the peptides identified in each phase-2 specimen, one tree per biological replicate (S-Trap; three injections pooled per specimen): (A) MNRJ 24305 and (B) MNRJ 24314, kept frozen at −20 °C; (C) MNRJ 24317 and (D) MNRJ 24325, stored in 80% ethanol at 4 °C. Each distinct non-contaminant peptide is placed at the lowest common ancestor of all database proteins that contain it, in the style of Unipept. Ribbon width and circle size scale with the square root of the number of peptides in each subtree; in each circle, the dark wedge is the share of the subtree’s peptides whose LCA is that node, and terminal nodes are fully filled. Colours: Eukaryota, blue; Bacteria, orange; Archaea, green; grey, the root (Organism), whose wedge marks peptides shared across domains. The percentages at top right give the share of each specimen’s peptides in Porifera, Bacteria and Poribacteria. The pooled phase-1 tree is in Supporting Figure S6.

Specimen-specific LCA trees had the same overall structure (Figure 9). Host assignments included Demospongiae, Hexactinellida and Calcarea, and symbiont assignments included Gammaproteobacteria, Alphaproteobacteria, Planctomycetia, Phycisphaerae, Nitrospiria, Actinomycetes, Flavobacteriia and the ammonia-oxidizing Nitrososphaeria in all four specimens. As in phase 1, assignments to several sponge classes reflect the distribution of matching sequences in the database and are not evidence that several sponge classes were present. Taxa not detected in the ethanol-stored specimens were represented by few peptides in the frozen ones: Homoscleromorpha (three peptides in each frozen specimen) was absent from both ethanol-stored specimens, and Brocadiia (five to six peptides) from MNRJ 24317. These absences are compatible with the lower depth and do not by themselves show community differences. Nitrospiria were also proportionally less represented in the ethanol-stored specimens (0.6–1.0% versus 2.0–2.1% of peptides).

A core of 199 protein groups was detected in every specimen (100 sponge and 99 symbiont groups), and 439 in at least three (Supporting Table S7 and Figure S5B). The symbiont core includes the Poribacteria SprA and MotA/TolQ/ExbB envelope proteins, chaperonins, glutamine synthetase, the P-II nitrogen regulator and the methanol dehydrogenase XoxF, and the host core includes cytoskeletal proteins. Their recurrent detection extends the Poribacteria-rich, nitrifying holobiont profile beyond the two specimens of phase 1.

#### Removing *Vibrio* from the database improved identifications without loss of taxa

*Vibrio* entries, culture isolates from other sponges, made up 5.4% of the residues of DB-2 but explained 0.1% of the PSMs in the 12 phase-2 runs (Supporting Table S10). Removing them and updating the contaminant list (DB-3) raised FragPipe identifications summed over the 12 runs: non-contaminant PSMs rose from 39,847 to 40,812 (+2.4%), peptides from 21,081 to 22,075 (+4.7%) and proteins from 11,992 to 12,740 (+6.2%). The gain was larger in the ethanol-stored specimens (+8.0% and +4.1% PSMs for MNRJ 24317 and MNRJ 24325) than in the frozen ones (+2.6% and +0.3%). Removing the Planctomycetota as well (DB-3a) would have doubled the gain (+6.0% PSMs, +9.2% peptides), but it would have removed a symbiont phylum present in every specimen. The entrapment test showed that the gain did not come at the cost of error: across the four tested injections, the estimated false discovery proportion fell from 1.9% with DB-2 to 1.5% with DB-3. With both databases the estimate exceeded the nominal 1%, so the FDR reported by the Sage search with global rescoring is slightly optimistic. In the LEMTEN pipeline, the larger contaminant list moved spectra from sample to contaminant proteins (contaminant spectra 3,421 to 3,867), and non-contaminant PSMs changed little (29,866 to 29,521).

### Implications for Proteomics of Non-Model Holobionts

The complementary analyses of the same 42 phase-1 runs support three methodological conclusions, and the phase-2 analysis of four specimens a fourth. Together they address workflow selection, the contribution of homology-driven identification, the comparison of pipelines and the influence of specimen history on proteome coverage.

#### Sample preparation determines depth more than search strategy does

Both pipelines ranked the runs almost identically (ρ = 0.98) and placed BST and A-PPT ahead of A-UT. The roughly twofold difference in acquired MS/MS spectra between A-UT and the other workflows was matched by a similar difference in identifications. The relative performance of the workflows was therefore robust to the analysis pipeline, although the contribution of individual preparation steps cannot be isolated from a comparison of complete workflows.

For sponge tissue rich in matrix components and symbionts, the results favour workflows that remove detergents and chaotropes before LC-MS/MS: S-Trap does this on a column and A-PPT by precipitation. S-Trap showed the smallest difference in depth between specimens (8%), whereas A-PPT differed by 39%, possibly through specimen-dependent co-precipitation of matrix components. We therefore recommend S-Trap as the default workflow for *V. ophiraphidites* under the conditions examined, with GeLC-MS/MS as a complement for selected large bacterial envelope proteins. Phase 2 supports this choice: S-Trap injections of the same digest varied by a median 22–25%, and a similar holobiont composition was recovered in all four specimens. Because each phase-2 specimen had a single preparation, however, this analysis does not establish extraction reproducibility or separate preparation effects from differences between specimens.

#### A conventional search of public databases leaves about a fifth of the interpretable evidence unused

The homology-driven layers increased assigned non-contaminant spectra by 21.3% over the conventional DB-1 search. The additional assignments came from the proteome of the closest sequenced relative, from MS-BLAST-style alignment of de novo tags rescored by precursor mass, and from an error-tolerant interpretation of the tag flanks. Most of the gain was host evidence, reflecting the divergence between *V. ophiraphidites* and the sponges represented in UniProt.

The mass-shift panorama shows what is lost without these layers. Thousands of spectra differ from their best database peptide by shifts consistent with single amino-acid substitutions such as T→V, A→G and N→E, with almost no decoy hits (4,571 targets against 47 decoys). For a species absent from UniProtKB, these substitutions are candidate sequence information not represented in the database. The homology-assigned peptides (Supporting Table S3) could guide a species-specific database once a genome or transcriptome of *V. ophiraphidites* becomes available.

Several observations support the additional homology-based assignments. Where both searches explained a spectrum, they agreed on 99.5% of peptides. Semi-specific explanations were 87 times more frequent among targets than decoys. The observed error among positive-control spectra matched the nominal 1% (1.0%), after correction of a decoy-reconstruction error that had doubled it in an earlier version of the pipeline. Agreement on shared spectra does not, however, validate every assignment obtained only by homology or error-tolerant searching.

#### Pipelines should be compared on equal databases and read as complementary

In the two scan-level comparisons against the same sequences, FragPipe identified about 1.4–1.5 times as many spectra as our Sage-based search. This difference reflects the complete search-and-rescoring configurations, including MSFragger fragment indexing and MSBooster-predicted spectra in Percolator^20^, rather than isolating the contribution of any single component. Our pipeline, in turn, contributes evidence that a fixed-database search cannot provide. For non-model organisms, a practical strategy is therefore to run the most sensitive conventional search available and add homology and error-tolerant tag layers on top of it. Matching databases is essential when pipelines are benchmarked, because database composition affects both identification depth and protein inference (Supporting Information, same-database comparison).

#### Specimen handling sets the ceiling for depth

With the same workflow, the same peptide loading by Qubit and a calibrated instrument, specimens stored long-term in ethanol at 4 °C gave about a quarter of the evidence of frozen ones and a larger share of contaminant spectra. These differences cannot be attributed specifically to preservation, because collection site and storage history were confounded. Protein yield per milligram of tissue, the contaminant share and the share of cysteine-containing PSMs in a search without fixed carbamidomethylation are nevertheless simple checks that flag such samples before a comparison is built on them.

## Limitations

Phase 1 included only two specimens, so differences between workflows are descriptive. In phase 2, each specimen had a single preparation, so specimen and preparation effects cannot be separated, and preservation was confounded with collection site, storage history and alkylation, which was not effective in the two frozen specimens for reasons that could not be established. BGEL likewise had a single preparation per specimen. Technical reinjections show measurement repeatability but do not add independent biological observations. Protein-group counts depend on the inference rule, and both pipelines report many single-peptide groups, as is typical for metaproteomes. Taxonomic assignment is constrained by shared sequences and database representation, and is reliable below the class level only for divergent proteins. An entrapment test estimated the false discovery proportion of the Sage-based database search at 1.5–1.9% for a nominal 1%. The homology layer is estimated per run, and with few accepted spectra per run its contribution varied widely between injections of the same digest; estimating its FDR over all runs of a preparation would stabilise it. Finally, homology-assigned peptides need confirmation against a *V. ophiraphidites* genome or transcriptome before they are used as sequence annotations, particularly where substitutions or unexplained mass shifts admit alternative interpretations.

## CONCLUSIONS

In phase 1, S-Trap digestion gave the deepest and most specimen-independent proteome of *Viles ophiraphidites*, with about 2,100 protein groups in each specimen and the lowest between-specimen variation among the workflows tested. Acetone precipitation reached similar depth in one specimen only, urea/thiourea in-solution digestion recovered about half as much, and GeLC-MS/MS complemented S-Trap for bacterial envelope proteins. Two independent search strategies supported this ranking.

Three homology-driven layers, the proteome of the closest sequenced relative available (*Halichondria panicea*), de novo tags aligned by MS-BLAST-style homology search and error-tolerant sequence-tag interpretation, raised the assigned evidence by 21.3% over a conventional search of public databases. They recovered sequence-divergent and semi-specific peptides that a fixed-database search does not report. Together, these workflows and searches provide a first proteomic characterization of this sponge holobiont, in which a host rich in cytoskeletal, chaperone and pentose-phosphate proteins coexists with a Poribacteria-dominated microbiome expressing cell-surface machinery and proteins consistent with nitrite oxidation by *Nitrospira*. In phase 2, the S-Trap analysis of four specimens recovered a similar composition and a core of 199 protein groups detected in all of them, about half from symbionts. Proteome depth followed specimen history and protein yield, although preservation was confounded with collection site and storage history, and reduction and alkylation were not effective in the two frozen specimens. Under the conditions examined, we recommend frozen storage (−20 °C) of sponge tissue intended for proteomics, and a check of protein yield and contaminant share before specimens are compared.

## ASSOCIATED CONTENT

### Supporting Information

The Supporting Information is available with this preprint.

- **Table S1.** Phase 1 per-run identification statistics, both pipelines (XLSX).
- **Table S2.** Phase 1 protein groups with source (Sage, homology or both), origin, taxon and Top3 abundance per run (XLSX).
- **Table S3.** Homology- and sequence-tag-assigned peptides with database homologue, precursor mass difference and annotation (XLSX).
- **Table S4.** Recurrent precursor mass shifts, target and decoy counts, annotation (XLSX).
- **Table S5.** Same-database comparison of FragPipe and the LEMTEN pipeline: identifications from *H. panicea*-only sequences excluded, and Sage re-searched on DB-1 for a subset of runs (XLSX).
- **Table S6.** Phase 2 specimens (collection site, storage, protein and peptide yield) and per-run statistics (XLSX).
- **Table S7.** Phase 2 protein groups per specimen, with core-proteome flag, spectral counts and Top3 abundance per run (XLSX).
- **Table S8.** Phase 2 peptide-level taxonomy (lowest common ancestor) per specimen (XLSX).
- **Table S9.** Cysteine forms in the 54 runs (alkylation check): PSMs with cysteine and the share of carbamidomethylated, disulfide-bonded, free, oxidized and other cysteines per run (XLSX).
- **Table S10.** Database evaluation: composition of DB-2 by taxon (residues and PSMs), *Geodia barretti* entries reclassified as bacterial, entrapment test of DB-2, DB-3a and DB-3, and FragPipe and LEMTEN identifications per phase-2 run with each database (XLSX).
- **Figures S1–S6.** De novo calibration and validation of the homology search (S1); phase 1 per-run QC (S2); duplicate-extraction correlations and PCA of the 42 runs (S3); heatmap of the 47 protein groups differing between workflows (S4); phase 2 homology gain, specimen overlap, PCA and injection reproducibility (S5); phase 1 peptide LCA tree, 42 runs pooled (S6) (PDF).

Files: Supporting_Information_Figures_S1-S6.pdf; Table_S1_per_run_statistics.xlsx; Table_S2_protein_groups.xlsx; Table_S3_homology_tag_spectra.xlsx; Table_S4_mass_shifts.xlsx; Table_S5_same_database_comparison.xlsx; Table_S6_quadruplicate_runs.xlsx; Table_S7_quadruplicate_protein_groups.xlsx; Table_S8_quadruplicate_taxonomy.xlsx; Table_S9_cysteine_forms.xlsx; Table_S10_database_evaluation.xlsx.

## Data Availability

The mass spectrometry proteomics data have been deposited to the ProteomeXchange Consortium via the PRIDE partner repository^33^ with the dataset identifier PXD079594, which will be made public upon journal publication. The dataset includes the raw files of both phases (phase 2: QE_018062–QE_018076). The LEMTEN pipeline, the search databases (DB-1, DB-2 and DB-3, with the list of relabelled *Geodia* entries) and analysis scripts will be made available at [repository URL].

## AUTHOR INFORMATION

### Corresponding Author

Magno Junqueira, LEMTEN, Instituto de Química, Universidade Federal do Rio de Janeiro, Rio de Janeiro, RJ, Brazil.

## Author Contributions

- C.S.A.: sample preparation, methodology, investigation, nLC-MS/MS, FragPipe analysis and writing.
- A.C.S.R.: sample preparation.
- G.R.S.: sample preparation.
- F.C.S.N.: nLC-MS/MS infrastructure and discussion.
- A.L.V.: natural products context and discussion.
- M.J.: conception, pipeline development, reanalysis, writing and supervision. [All authors: confirm.]

## Notes

The authors declare no competing financial interest.

## ACKNOWLEDGMENTS

**In memoriam.** This work is dedicated to Prof. Guilherme Ramos da Silva Muricy (1964–2026), full professor in the Departamento de Invertebrados of the Museu Nacional, Universidade Federal do Rio de Janeiro, who died on 19 August 2026, aged 62. Trained in Marine Biology at UFRJ, in Oceanography at the Université d’Aix-Marseille and in Biochemistry at UFRJ, he reactivated the Laboratório de Porifera of the Museu Nacional as an undergraduate in 1986. He built it into one of Brazil’s leading sponge research groups and published more than a hundred works on sponge taxonomy and chemistry. His team collected and identified the specimens studied here, then as *Topsentia ophiraphidites*, and without his decades of work on Brazilian sponges this study would not exist. We extend our deepest condolences to his family, colleagues and students (UFRJ, 21 August 2026).

We also thank Joana Sandes and the team of the Laboratório de Porifera, Museu Nacional/UFRJ, for specimen collection during the 2022 Fernando de Noronha expedition. This work was supported by CNPq, FAPERJ and CAPES [grant numbers]. Parts of the analysis code and of the manuscript text were drafted with the assistance of generative AI (ChatGPT, OpenAI; Claude, Anthropic) and were reviewed and verified by the authors.

## ABBREVIATIONS

A-UT: urea/thiourea in-solution digestion
A-PPT: acetone precipitation
BST: S-Trap digestion
BGEL: GeLC-MS/MS
DB-1: laboratory database
DB-2: DB-1 plus *Halichondria panicea* proteome
DB-3: DB-2 without *Vibrio* entries and with an updated contaminant list
FDR: false discovery rate
PSM: peptide-spectrum match
SAF: spectral abundance factor
XIC: extracted-ion chromatogram.

