## Supplementary material for "Sample Preparation and Homology-Driven Search Shape the First Holobiont Proteome of the Marine Sponge *Viles ophiraphidites* (formerly *Topsentia ophiraphidites*)": Supporting_Information_Figures_S1-S6.pdf

**Contents**

Figure S1. De novo sequencing calibration and validation of the homology search

Figure S2. Per-run quality control

Figure S3. Reproducibility between extractions and PCA of all runs

Figure S4. Protein groups differing between workflows

Figure S5. Phase 2: biological quadruplicate with S-Trap

Figure S6. Phase 1 peptide-level taxonomy (LCA tree, 42 runs pooled)

**Supporting tables (separate XLSX files):**

Table S1. Per-run identification statistics, both pipelines

Table S2. Protein groups with origin, spectral counts and Top3 abundance per run

Table S3. Homology- and sequence-tag-assigned spectra

Table S4. Recurrent precursor mass shifts and annotation

Table S5. Same-database comparison of FragPipe and the LEMTEN pipeline

Table S6. Phase 2: specimens (site, storage, protein and peptide yield) and per-run statistics

Table S7. Phase 2: protein groups per specimen, with core proteome flag

Table S8. Phase 2: peptide-level taxonomy (LCA) per specimen

Table S9. Cysteine forms in the 54 runs (alkylation check)

Table S10. Database evaluation (DB-2, DB-3a, DB-3)

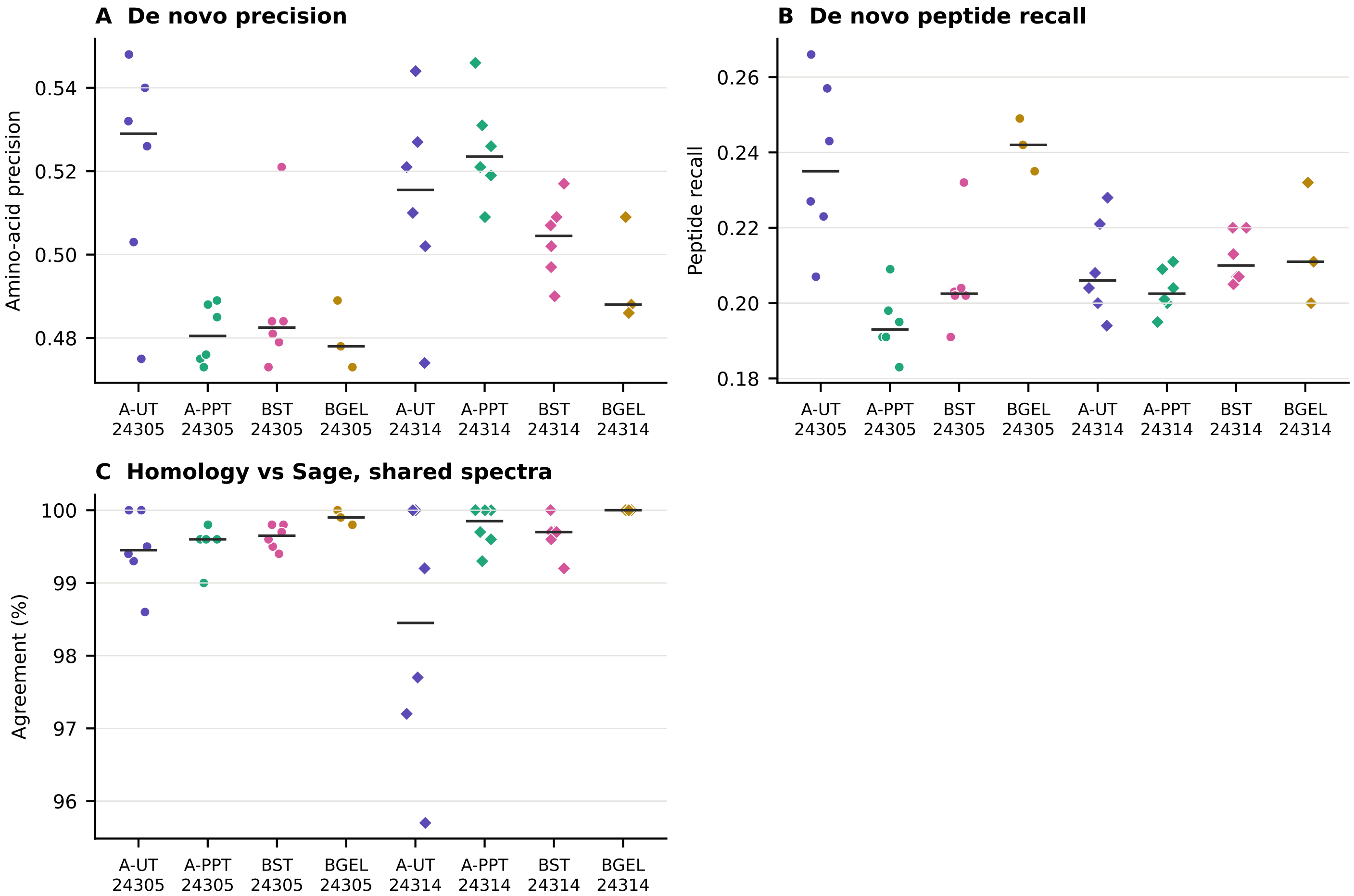


Figure S1. De novo sequencing calibration and validation of the homology search. (A) Amino-acid precision and (B) peptide recall of de novo calls measured against the 1% FDR Sage PSMs of the same run. (C) Percentage of spectra identified by both the homology search and Sage that received the same peptide sequence (I = L). Points are runs (circles MNRJ 24305, diamonds MNRJ 24314); bars are medians.

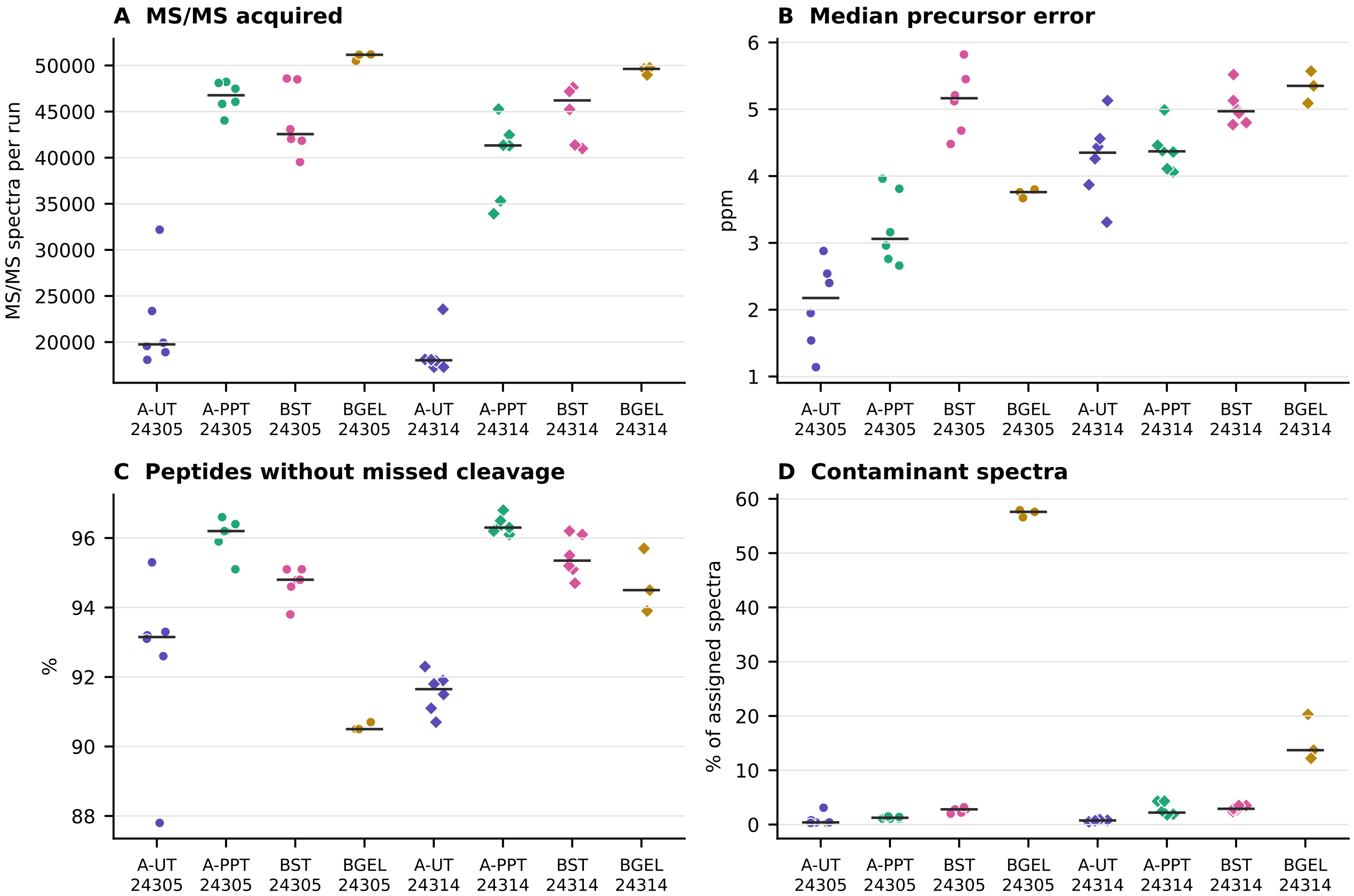


Figure S2. Per-run quality control (LEMTEN pipeline). (A) MS/MS spectra acquired per run; (B) median precursor mass error of 1% FDR PSMs; (C) share of peptides without missed cleavages; (D) share of assigned spectra matching only contaminants (keratins, trypsin), high in the gel workflow.

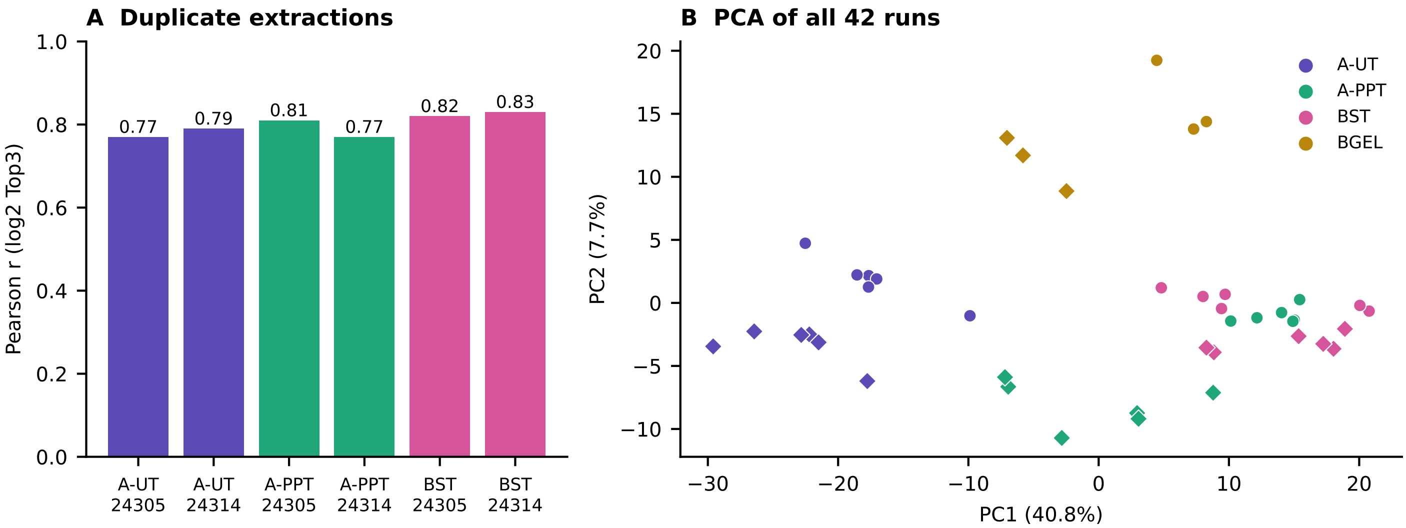


Figure S3. Reproducibility. (A) Pearson correlation of log2 Top3 abundance between duplicate extractions (injection means); BGEL had one preparation per specimen. (B) PCA of the 42 runs (log2 Top3, groups quantified in all runs); circles MNRJ 24305, diamonds MNRJ 24314.

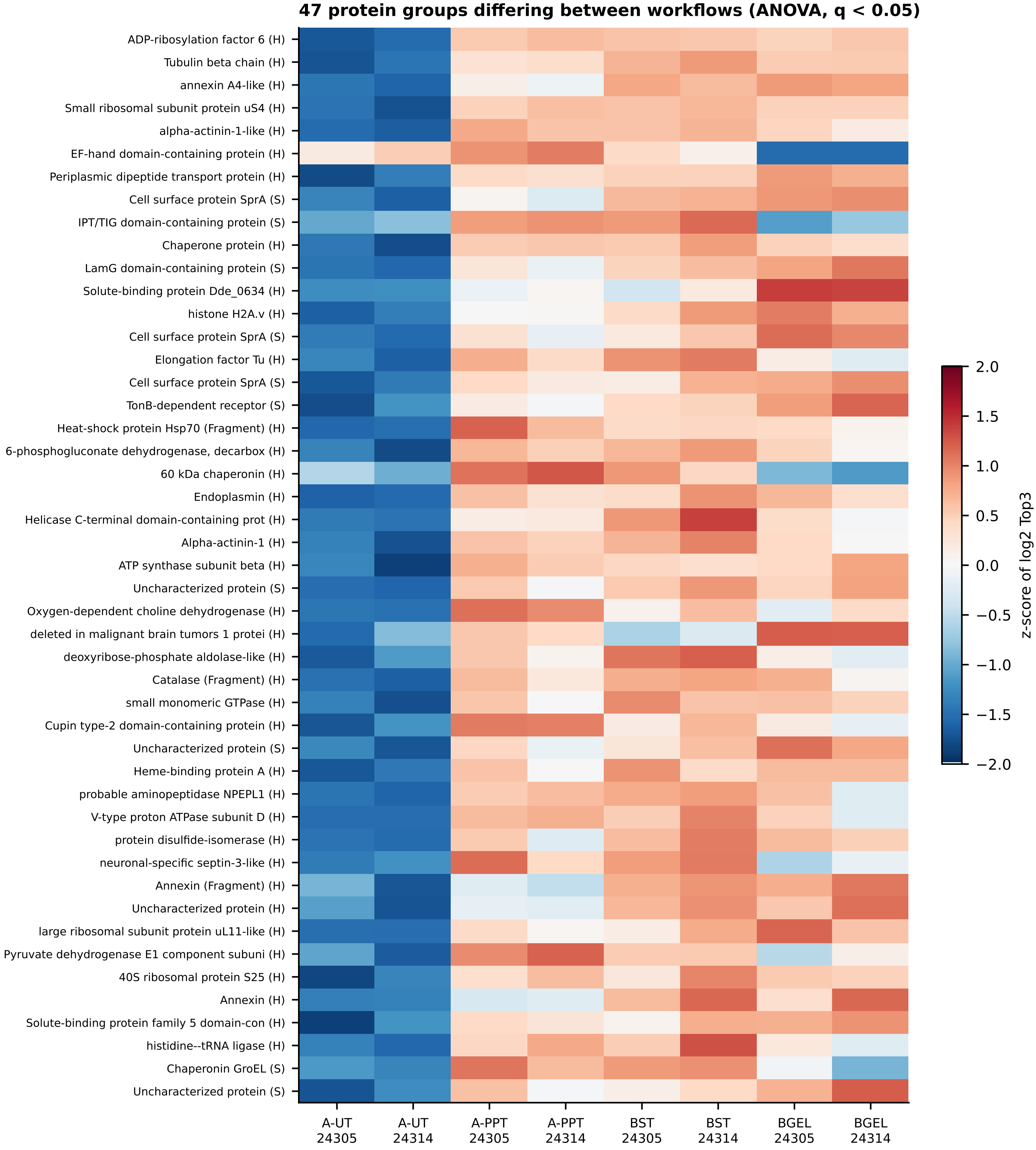


Figure S4. Protein groups whose abundance differed between workflows (one-way ANOVA on specimen means of log2 Top3, Benjamini-Hochberg q < 0.05, among 268 groups quantified in all workflows). H, host (sponge); S, symbiont.

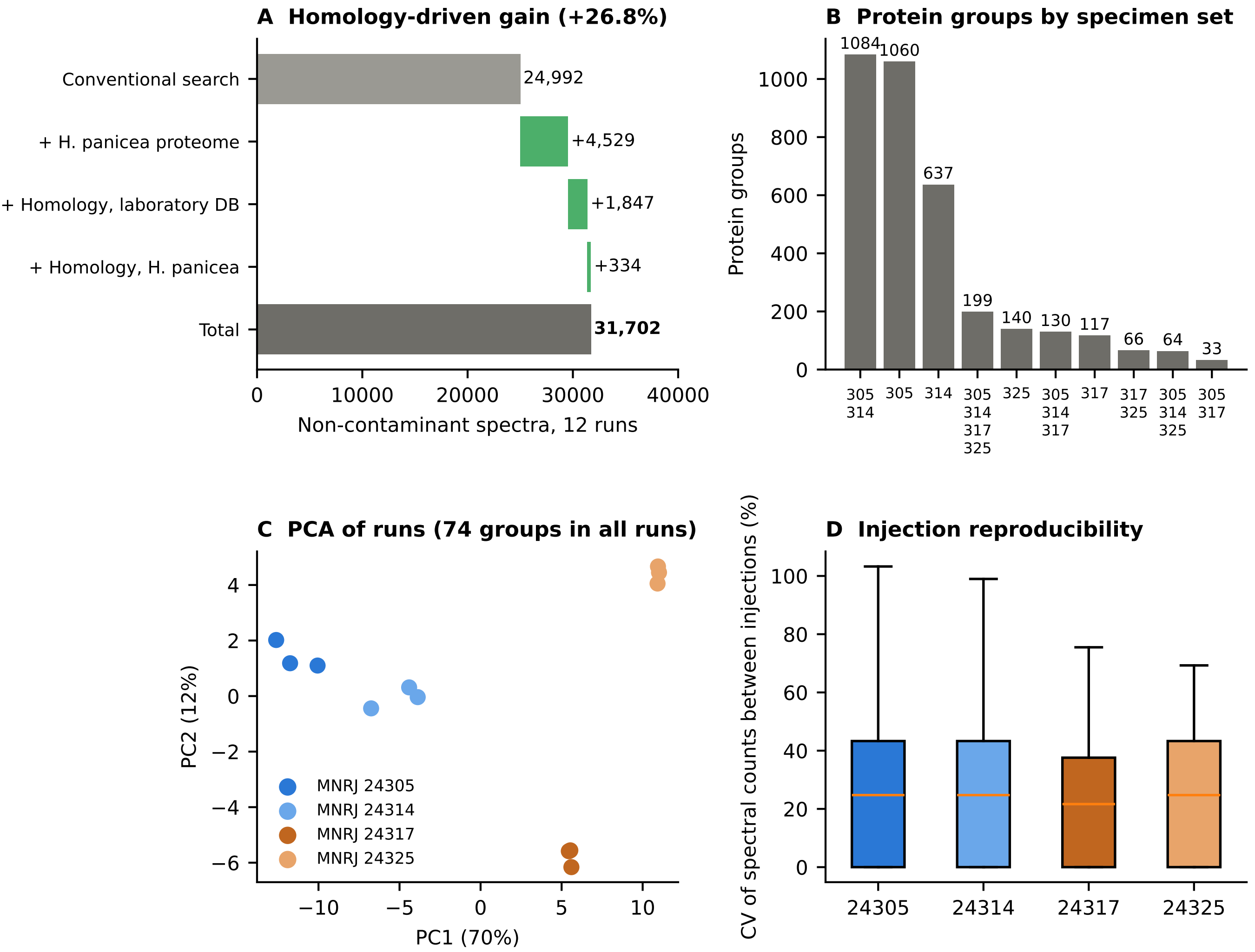


Figure S5. Phase 2 biological quadruplicate (S-Trap; MNRJ 24305 and 24314 frozen at −20 °C, MNRJ 24317 and 24325 stored in ethanol at 4 °C; one preparation each, three injections). (A) Spectra added by each homology-driven layer over a conventional search, 12 runs, 1% FDR. (B) Protein groups (contaminants excluded) by the set of specimens in which they were detected; 199 groups were found in all four. (C) PCA of the 12 runs (log2 Top3, median-centred, groups quantified in all runs). (D) Coefficient of variation of spectral counts between the three injections of each specimen (groups detected in all three).

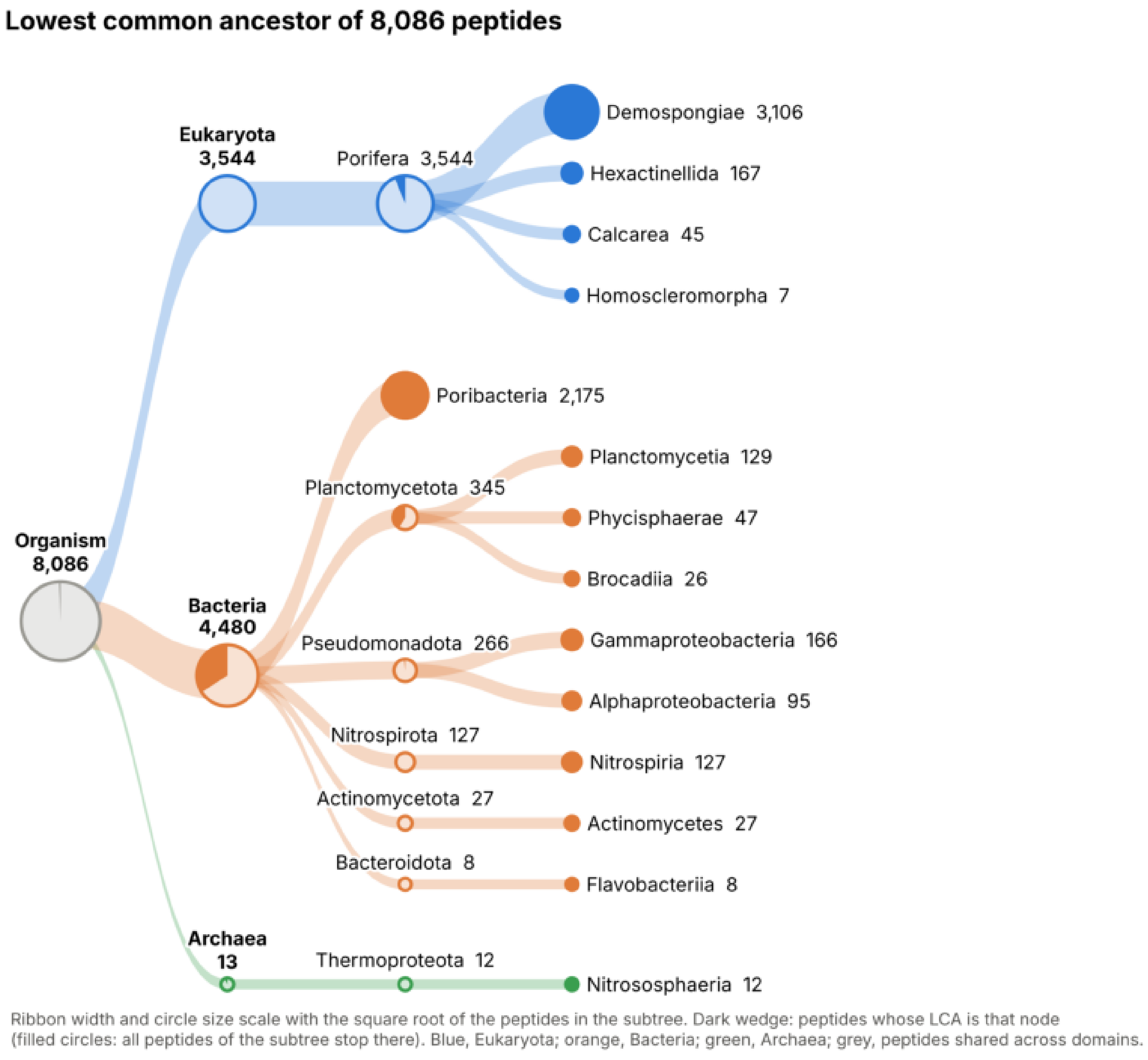


Figure S6. Phase 1 peptide-level taxonomy (both specimens, all 42 runs pooled). Each of the 8,086 distinct non-contaminant peptides is placed at the lowest common ancestor of all database proteins that contain it, in the style of Unipept. Ribbon width and circle size scale with the square root of the number of peptides in each subtree. In each circle, the dark wedge is the share of the subtree's peptides whose LCA is that node; terminal nodes are fully filled. Colours: Eukaryota, blue; Bacteria, orange; Archaea, green; grey, the root (Organism), whose wedge marks the 49 peptides shared across domains. Entries of the Geodia barretti proteome whose sequences are bacterial were relabelled as unclassified Bacteria (Methods). Per-specimen trees from phase 2 are in Figure 9 of the main text.
